# Healthy-to-Stroke Translation of EEG-Based BMIs: EEG Characterization and Reinforcement Learning–Based Decoder Evaluation

**DOI:** 10.64898/2026.06.23.733831

**Authors:** Zander Via, Ava Kruse, Bhoj Raj Thapa, Jihye Bae

## Abstract

**Purpose:** EEG-based brain–machine interfaces (BMIs) may support assistive technologies for individuals with stroke-related motor impairment by translating neural activity into control commands for external devices. However, post-stroke neural reorganization and interindividual EEG variability challenge reliable decoding. This study characterized motor imagery EEG features in healthy and acute stroke participants and evaluated whether population-trained Q-learning Kernel Temporal Difference (Q-KTD) decoders could improve individual stroke decoding through transfer learning. These analyses assess the feasibility of healthy-to-stroke translation for EEG-based BMI neural decoding.

**Materials and Methods:** Publicly available motor imagery EEG datasets from healthy participants (n = 109) and individuals with acute stroke (n = 50) were analyzed using left- and right-hand motor imagery trials. The datasets were selected because of their relatively large sample sizes and comparable motor imagery tasks. EEG characterization included baseline and motor imagery-period band power, ERD/ERS, hemispheric asymmetry, and time–frequency representations. For Q-learning Kernel Temporal Difference (Q-KTD) decoding, filtered time-domain EEG from 0– 0.5 s after motor imagery onset was used as the neural-state input. A Q-KTD model trained on the healthy population was transferred to individual stroke participants, and repeated Monte Carlo simulations compared decoding performance with and without transfer learning across multiple learning epochs.

**Results:** Healthy and acute stroke participants showed shared motor imagery–related EEG structure, including post-onset mu-band suppression, while the stroke group exhibited greater interparticipant variability, more diffuse time– frequency modulation, and altered hemispheric asymmetry. No channel-level healthy–stroke differences in windowed band power remained significant after false discovery rate correction. Healthy-source transfer learning improved first-epoch Q-KTD success rates in 29 of 50 stroke participants (58%). Across all participants, mean success rate increased from 49.46% without transfer learning to 51.82% with transfer learning. Among participants showing positive transfer, the mean gain was 7.34% and the maximum gain was 18.75%. However, 21 participants showed negative transfer, demonstrating substantial subject-level variability.

**Conclusion:** Healthy-source Q-KTD transfer learning improved first-epoch motor imagery BMI decoding for a majority of acute stroke participants, supporting the offline feasibility of population-informed Q-KTD decoding in stroke. These early performance gains may reduce subject-specific calibration burden, although substantial interparticipant variability and negative transfer indicate the need for individualized transfer-selection or adaptation strategies.

**Assistive Technology Implications:**

- EEG-based brain–machine interfaces may support assistive technologies for individuals with stroke-related motor impairment by translating motor imagery–related neural activity into control commands for external devices.
- Healthy-to-stroke transfer learning may improve early BMI neural-decoder performance and potentially reduce the amount of subject-specific calibration required.
- The findings support the offline feasibility of Q-KTD for motor imagery BMI neural decoding in individuals with acute stroke.
- Substantial interparticipant variability and negative transfer suggest that individualized source-model selection or adaptation strategies may be needed for reliable post-stroke BMI implementation.
- Physiological EEG characteristics, including ERD/ERS and hemispheric asymmetry, may provide candidate markers for future transfer-selection strategies, although their predictive value requires direct validation.

## Introduction

Electroencephalography (EEG) provides a practical neural sensing approach for brain-machine interface (BMI) development because it can measure time-varying neural activity non-invasively from the scalp. Compared with invasive neural recordings, EEG has lower spatial resolution, but it offers high temporal resolution, lower implementation burden, portability, and compatibility with repeated use across different environments. These characteristics make EEG well suited for BMIs that translate neural activity into control commands for external systems, including computer cursors, robotic devices, neuroprosthetic interfaces, and other assistive technologies (1–5). The practical challenge is that EEG is not only a sensing modality. It is also a highly variable signal source that requires decoding methods capable of adapting across users and conditions.

This variability is one of the major barriers to reliable EEG-based BMI use. EEG signals are weak, artifact-sensitive, nonstationary within users, and variable across subjects, sessions, devices, and tasks (6,7). As a result, a decoder trained under one condition may not perform reliably when applied to another user or population. This problem becomes especially important when BMI systems are developed for individuals affected by stroke, where the target population may share some movement-related neural structure with healthy participants while also showing neurological changes that alter how this structure is expressed.

Stroke can disrupt the neural systems involved in movement planning, motor execution, and sensorimotor integration. These effects are not limited to the injured region alone. After stroke, motor-related activity may be influenced by changes in cortical excitability, corticospinal pathway integrity, communication between hemispheres, recruitment of ipsilesional and contralesional regions, and reorganization of functional motor networks (8–13). These changes can affect how movement-related information is represented in the brain, even when a stroke participant and a neurologically intact participant are performing the same task.

For EEG-based BMI development, a key factor is that stroke EEG may contain both shared and altered structure. Shared structure matters because it creates the possibility that information learned from healthy participants can help initialize a decoder for stroke participants. Altered structure matters because it explains why a fixed model trained only on healthy data may fail or provide inconsistent benefit. Stroke participants may show movement-related EEG patterns that overlap with those observed in healthy participants, while also showing changes in timing, magnitude, spatial distribution, hemispheric balance, and trial-to-trial consistency (10–17). Quantitative EEG studies also suggest that stroke can affect broader spectral organization, including increased low-frequency activity, hemispheric asymmetry, and attenuation of faster rhythms (14–17).

Motor imagery (MI) was used in this study as the task paradigm for eliciting movement-related EEG activity. MI involves imagining movement without overt execution and is widely used in EEG-based BMI research because it can produce measurable changes over sensorimotor scalp regions (18–23). In this manuscript, MI is not treated as the primary endpoint. Instead, it provides a controlled task structure for comparing EEG organization between healthy and stroke participants and for evaluating whether source-population EEG information can improve BMI decoding. This distinction is important because the broader goal is to understand whether movement-related EEG features can support decoder adaptation for individuals affected by stroke, not to characterize MI physiology in isolation.

The most established EEG features used in MI-based BMI studies are changes in sensorimotor rhythms, especially mu and beta activity. During imagined or attempted movement, these rhythms often decrease in power relative to baseline, a response known as event-related desynchronization. Power may also increase after task completion as event-related synchronization or beta rebound (19–23). These features provide interpretable markers for movement-related decoding. In healthy participants, left-hand and right-hand MI often produce organized differences over central electrodes such as C3 and C4, which serve as practical scalp-level markers of hemispheric sensorimotor organization (20–23). In stroke participants, similar sensorimotor modulation may be present, but it may appear weaker, delayed, spatially shifted, more bilateral, or more variable (10–13). For this reason, sensorimotor rhythm modulation and C3-C4 balance were used to examine both shared movement-related EEG structure and stroke-related deviations that may affect decoder transfer.

A broader spectral analysis was also included because the EEG differences relevant to BMI adaptation may not be limited to conventional mu and beta rhythms. Low-frequency EEG activity can reflect spectral slowing, cortical-state changes, or movement-preparatory activity (14–17,24–27), while higher-frequency activity may provide additional information related to movement or BMI performance (28–32). Analyzing EEG activity from 0.1 to 80 Hz allows the study to examine whether healthy and stroke participants differ only in conventional sensorimotor rhythms or across a broader EEG profile. This matters because a BMI decoder may be influenced by both task-related sensorimotor modulation and background differences in neural signal organization.

Transfer learning provides one approach for reducing calibration demands in EEG-based BMI systems. Rather than training a decoder only from the target user’s data, transfer learning uses information learned from a source subject or source population to improve learning in a target user (7,33,34). Shared EEG structure can provide a useful starting point for decoder initialization, while target-user adaptation can account for features that differ from the source population. In this study, healthy-to-stroke transfer tested whether EEG structure learned from neurologically intact participants could improve decoding in stroke participants.

Q-learning Kernel Temporal Difference (Q-KTD) was selected as the decoding framework because EEG-based BMI control requires a decoder that can learn from noisy, high-dimensional, and subject-specific neural signals while adapting through feedback. In a conventional static classifier, EEG features are mapped to intended movement classes using a fixed supervised model; however, this approach does not directly represent the reward-driven interaction between the BMI decoder, the user, and the task environment. In an RLBMI formulation, the EEG feature vector at each trial is treated as the state, the decoder-selected movement direction is treated as the action, and task performance, such as whether the cursor reaches the correct target, is used to assign reward (35–39). Q-learning is appropriate for this setting because it estimates the action-value function, or the expected cumulative reward for selecting a particular action in a given neural state, and learns policies that maximize long-term reward. Kernel Temporal Difference learning further extends this framework by using kernel-based nonlinear function approximation, allowing the decoder to model complex relationships in EEG feature space without requiring a tabular representation of all possible states (36–42). This makes Q-KTD an appropriate framework for evaluating whether source-model initialization can improve early decoder adaptation when the target user is a stroke participant.

This study addresses whether shared EEG structure across healthy and stroke populations can be used to reduce early decoder calibration demands without ignoring stroke-related neural heterogeneity. The primary aim was to determine whether transfer learning improves first-epoch Q-KTD decoding performance in stroke participants. The secondary aim was to characterize both similarities and differences between healthy and stroke EEG across spectral, temporal, spatial, and hemispheric features. We hypothesized that transfer learning would improve early decoder performance for many stroke participants because movement-related EEG structure may be partially shared across populations. However, we also expected the magnitude of benefit to vary across individuals because EEG organization after stroke is heterogeneous. Together, these analyses evaluate whether source-population EEG structure can support adaptive BMI decoder initialization for individuals affected by stroke.

## Materials and Methods

The aim of this study was twofold: first, to extract and analyze EEG features, and second, to implement those features within the Q-KTD decoding framework. The initial analysis examined whether acute stroke participants exhibited distinct EEG feature patterns compared with neurologically intact participants during binary left- and right-hand motor imagery trials. The extracted filtered EEG features were then formatted for use as inputs to the Q-KTD neural decoder. Together, this design addressed two linked goals: evaluating whether stroke-related motor imagery EEG differs from healthy motor imagery EEG, and generating consistent, decoder-ready EEG inputs for subsequent Q-KTD analysis.

### EEG datasets

Two publicly available EEG datasets were analyzed: one dataset from neurologically intact participants and one dataset from acute stroke participants. The healthy dataset was used as the neurologically intact comparison dataset, and the stroke dataset was used as the for analysis of EEG patterns in individuals affected by stroke as well as subjected to transfer learning. Dataset-specific acquisition details are summarized in the next two paragraphs, while task alignment, channel harmonization, preprocessing, and feature extraction are described in the following sections.

Healthy EEG data were obtained from the PhysioNet EEG Motor Movement/Imagery Dataset (43). This dataset was selected as the neurologically intact source population because it provides a large, standardized collection of motor execution and motor imagery EEG recordings from 109 healthy participants as well as keeping a similar experimental paradigm to the acute stroke dataset. The recordings were acquired using the BCI2000 instrumentation system (44) and include more than 1500 EEG trials across the full experimental protocol. EEG was recorded from 64 scalp electrodes during visually cued movement and imagery tasks, with a sampling frequency of 160 Hz. The PhysioNet recordings are provided as raw EEG without post-processing or re-referencing. The original acquisition used the right mastoid as ground and the right earlobe as reference, and no EOG channels were recorded. Participant age and sex information were not provided in the public repository and were therefore not included in demographic comparisons (45).

Each participant completed 14 experimental runs. Runs 1 and 2 were one-minute baseline recordings with eyes open and eyes closed, respectively. Runs 3 through 14 consisted of three two-minute repetitions of four task conditions: executed unilateral fist movement, imagined unilateral fist movement, executed bilateral fist or foot movement, and imagined bilateral fist or foot movement. For this study, only imagined unilateral fist movement runs, consisting of runs 4, 8, and 12, were used for the purposes of motor imagery analysis. During the unilateral fist tasks, a target appeared on either the left or right side of the screen, instructing the participant to either open and close the corresponding fist or imagine opening and closing the corresponding fist until the target disappeared, followed by relaxation (43).

Motor execution runs, bilateral hand/foot imagery runs, foot imagery trials, and baseline-only runs were excluded from the primary decoder-oriented analysis. In the selected unilateral imagery runs, EDF+ event annotations were used to identify task timing. The T0 annotation indicated rest, T1 indicated left-fist imagery, and T2 indicated right-fist imagery. Because the meanings of T1 and T2 differ across other PhysioNet task runs, event labels were interpreted according to run type before trial extraction (43).

Acute stroke EEG data were obtained from the dataset described by Liu et al. and deposited in Figshare (46,47). This dataset contains recordings from 50 acute stroke participants, including 39 males and 11 females aged 31 to 77 years, with time after stroke ranging from 1 to 30 days. Each participant completed 40 cue-guided motor imagery trials involving imagined left-hand or right-hand grasping of a spherical object. Each trial lasted 8s and included an instruction stage, a 4s motor imagery stage, and a break period. During the motor imagery stage, participants imagined grasping the object with the cued hand while a gripping-motion video was displayed. EEG was recorded using a wireless multichannel EEG acquisition system (ZhenTec NT1, Xi’an ZhenTec Intelligence Technology Co., Ltd., China) with electrodes arranged according to the international 10-10 system and sampled at 500 Hz. The reference electrode was positioned at CPz and the ground electrode was positioned at FPz (46,47). The raw files contain 33 channels: 30 EEG channels, 2 electrooculography (EOG) channels, and 1 marker event channel. The marker event channel provides task timing information, while the trial label identifies the task class, with “1” corresponding to left-hand motor imagery and “2” corresponding to right-hand motor imagery. The repository also includes stimulus files, which refer to the visual and instructional cue materials used during the left- and right-hand grasping tasks. For additional information on the data acquisition of this dataset, please refer to the original publication (46, 47).

### Task Alignment

The healthy and stroke datasets were aligned to a common binary left-hand versus right-hand motor imagery framework to match the intended decoder objective. In the healthy dataset, only unilateral left- and right-fist motor imagery runs were retained; baseline runs, motor execution runs, bilateral hand/foot tasks, and foot imagery tasks were excluded. In the stroke dataset, all labeled trials were retained because the task was already limited to cue-guided left- and right-hand grasping imagery. It should be noted that while the motor imagery tasks were similar across the datasets, recordings of the datasets differ. The healthy dataset consisted of a continuous recording whereas the stroke dataset consisted of trial-based and consistently timed events for recordings. They also differ via the healthy dataset alternating between rest periods and motor imagery between trials, whereas the stroke dataset starts trials with a two second period of instruction, followed by motor imagery, and then a rest period.

After alignment, both datasets represented the same decoder-level problem: distinguishing left-hand from right-hand motor imagery. However, the datasets still differed in acquisition system, sampling rate, channel montage, event timing, and imagined movement. These differences were addressed during preprocessing by removing non-consistently aligned channels, harmonizing both datasets to a shared EEG montage, resampling to a common sampling rate, applying the same filtering approach, and extracting dataset-specific motor imagery-aligned analysis windows. This allowed healthy and stroke EEG to be compared within a common left-versus-right motor imagery framework while acknowledging unavoidable cross-dataset differences.

### Preprocessing

All EEG preprocessing was performed in MATLAB R2025a. The preprocessing pipeline was designed to convert healthy and stroke datasets into a shared, subject-consistent format for feature extraction and Q-KTD decoder analysis. Healthy data were processed from EDF files, while stroke data were processed from subject-specific .mat files. The two datasets differed in acquisition system, sampling rate, channel montage, reference scheme, and task timing. Therefore, preprocessing standardized the data structure while preserving as much dataset-specific information as possible for the most full-scaled analysis possible.

### Re-referencing and Channel Harmonization

To ensure consistent analysis across datasets, channel harmonization was performed to preserve EEG channels consistent between the two datasets prior to feature extraction and decoder analysis. This is essential to retain as much information in each dataset as possible while ensuring fair analysis. For the healthy PhysioNet data, EEG channels were re-referenced to CPz before channel harmonization. The CPz time series was subtracted sample-by-sample from each EEG channel. After re-referencing, CPz was then removed from the shared EEG montage because the CPz-referenced CPz channel becomes zero-valued and does not provide usable physiological information. For the stroke dataset, electrodes that do not provide data consist with EEG, including electrooculography and marker/event channels, were removed before feature extraction.

Non-EEG channels, including EOG and event or trigger channels, as well as all reference channels were removed before feature extraction. After removal of these channels from the following analysis, the remaining EEG channels that were retained are as follows: FP1, FP2, FZ, F3, F4, F7, F8, FCZ, FC3, FC4, FT7, FT8, CZ, C3, C4, T3, T4, CP3, CP4, TP7, TP8, PZ, P3, P4, T5, T6, OZ, O1, and O2.

### Resampling

To ensure that filtering, epoch indexing, time-window definitions, and feature extraction were aligned and scaled for every subject, all data were analyzed at a common sampling rate of 160 Hz. Stroke recordings were downsampled from 500 Hz to 160 Hz. Healthy EDF recordings were also checked for their original sampling rate before filtering and epoching. If a healthy EDF file was not sampled at 160 Hz, the continuous EEG data were resampled to 160 Hz before filtering. 160 Hz was chosen as the sampling frequency as it preserved the most amount of frequency information as allowed between the datasets for analysis between low-delta oscillation and high-gamma activity.

### Filtering

EEG activity was isolated using zero-phase digital filtering in MATLAB with fourth-order Butterworth filters implemented using butter and filtfilt. Zero-phase filtering was used to reduce phase distortion and preserve temporal alignment between EEG activity, cue onset, and the feature-extraction windows. Filtered signals were generated for delta/slow-delta activity from 0.1–4 Hz, theta activity from 4–8 Hz, mu activity from 8–13 Hz, beta activity from 13– 30 Hz, and gamma activity above 30 Hz. Because all EEG signals were analyzed at 160 Hz, the highest theoretically analyzable frequency was 80 Hz according to the Nyquist limit; therefore, gamma-band analyses were interpreted within the valid 30–80 Hz frequency range.

The 0.1–4 Hz delta/slow-delta band was included because low-frequency EEG activity is strongly relevant in stroke. Quantitative EEG studies have shown that ischemic stroke is commonly associated with spectral slowing, increased low-frequency power, hemispheric asymmetry, and changes in indices such as the brain symmetry index and delta-related measures, which can provide information about ischemic injury and functional outcome (14–17). The lower cutoff of 0.1 Hz was selected to retain very slow EEG fluctuations and slow cortical activity that may reflect altered cortical state, preparatory activity, or post-stroke low-frequency abnormalities rather than removing these components with a more conventional 0.5 or 1 Hz high-pass threshold (26,27). Thus, the delta/slow-delta band was included to capture broad low-frequency changes that may differentiate acute stroke EEG from healthy EEG and may influence decoder transferability.

Theta activity from 4–8 Hz was included because stroke-related EEG slowing is not limited to the delta range and may extend into neighboring low-frequency bands. Theta-band activity may reflect altered cortical state, attentional demands, task preparation, and low-frequency changes associated with neurological injury (14–17,24–27). Including theta allowed the analysis to determine whether healthy-stroke differences and transfer-relevant similarities were restricted to conventional sensorimotor rhythms or were also present in broader low-frequency EEG organization.

Mu and beta bands were included because they are standard EEG frequency ranges for motor imagery-based BMI analysis. During imagined or attempted movement, sensorimotor rhythms in the mu and beta ranges often decrease in power relative to baseline, a response known as event-related desynchronization, and beta activity may later increase as event-related synchronization or beta rebound (19–23). These rhythms are commonly used for MI decoding and are often evaluated over sensorimotor scalp regions, including electrodes such as C3 and C4 (20–23). Therefore, mu and beta features were included as the primary motor-related EEG bands for evaluating shared and altered sensorimotor structure between healthy and stroke participants.

Gamma activity was included as an exploratory higher-frequency feature family because prior EEG and MEG studies suggest that gamma-band activity can contain movement-related or BMI-relevant information during motor imagery and may be associated with BCI performance (28–32). However, scalp-recorded gamma activity is more susceptible to muscle activity, high-frequency noise, and recording artifacts than lower-frequency sensorimotor rhythms. For this reason, gamma-band features were retained for exploratory comparison but interpreted cautiously relative to delta, theta, mu, and beta features.

### Epoching and Analysis Window

Healthy and stroke epochs were segmented and extracted using dataset-specific event timing information for the motor imagery event during the trial. For the healthy dataset, T1 and T2 EDF annotations were identified in the selected unilateral fist imagery runs to identify motor imagery onset. Healthy trials were extracted from –4.1 to +4.1 relative to motor imagery onset. This window was chosen because T0, T1, and T2 duration varied across healthy subjects. Therefore, aligning every healthy trial to motor imagery onset ensured that onset occurred at the same sample location across all trials. If a trial did not contain the full pre- or post-onset interval, missing samples were padded with NaN rather than zeros to avoid biasing averages or power estimates as well as to make the size of trials uniform so that MI onset is consistent across all trials.

For the stroke dataset, trials were segmented using the known 8s trial structure. The first 2 seconds represented instruction for the task, followed by 4 seconds of motor imagery, and then 2 seconds of rest. This ensured that motor imagery onset always occurred at the same sample location.

After epoching, specific features were extracted from each of the selected bands. Feature extraction used the baseline and motor imagery windows saved in each subject-level preprocessing file. The current analysis used both a 0.5 second and a 1 second analysis window. For healthy subjects, the baseline window was the analysis window size interval immediately before motor imagery onset, and the motor imagery window was the analysis window size interval immediately after motor imagery onset. For stroke subjects, the motor imagery window was the analysis window size interval immediately after the known cue onset, and the baseline/rest window was taken from the late-trial rest period happening between six and eight seconds during each trial. These windows were used consistently for band power, ERD/ERS, asymmetry, topographical/channel-power features, and statistical analysis.

### Trial Labels and Integrity Checks

Trial labels were retained after epoching so that each trial remained linked to its left- or right-hand motor imagery class. Filtered epochs were stored as channels by samples by trials for each frequency band. Valid-trial masks were compared across all filtered bands, and processing was stopped if trial inclusion differed between bands. This ensured that feature values across frequency bands were extracted from the same set of trials.

### Subject-Specific Output

For each participant, preprocessing saved a MATLAB structure containing the subject identifier, sampling rate, epoch time vector, channel labels, trial labels, filtered epoched EEG data, baseline window, motor imagery window, cue timing information, and relevant event metadata. Healthy outputs retained EDF annotation timing information, including motor imagery event timing and preceding rest durations. Stroke outputs retained the trial timing information from the original task structure. This common subject-level output allowed the same downstream feature extraction, visualization, statistical analysis, and decoder-preparation routines to be applied to both datasets.

### Features

Feature extraction was performed in MATLAB using the preprocessed subject-level files generated from the preprocessing pipeline. For each participant, the analysis script loaded the filtered EEG epochs, channel labels, time vector, trial labels, baseline window, and motor imagery window saved during preprocessing. Feature extraction was performed at the subject level before group averaging so that each participant, rather than each trial, served as the primary statistical unit.

For each subject, filtered EEG data were represented as a three-dimensional epoch array.

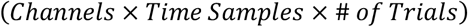

Subject-level feature values were then reduced into feature vectors and assembled into subject-by-feature matrices for statistical analysis.

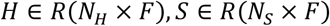

where *H* represents the healthy participant feature matrix, *S* represents the stroke participant feature matrix, *N*_*H*_ and *N*_*S*_ are the number of healthy and stroke participants, and *F* is the number of features for a given feature family. Each row represented one participant and each column represented one EEG feature.

The feature set was selected to characterize motor imagery EEG from complementary perspectives: cue-aligned time-domain responses, frequency-specific windowed power, task-related desynchronization/synchronization, hemispheric asymmetry, time-frequency modulation, and spatial channel-power organization. These features were used to determine whether acute stroke participants showed EEG patterns that differed from healthy participants and to identify feature families that may be useful for stroke-focused adaptive neural decoder development.

### Cue-Aligned Event-Related Potentials

Cue-aligned event-related potentials were computed to visualize time-domain EEG responses surrounding motor imagery onset. For each participant, filtered EEG epochs were averaged across trials separately for left-hand and right-hand motor imagery. ERP features were extracted across the full harmonized channel montage for both the broad motor imagery band and the delta/slow-delta band.

For each subject, channel, task class, and band, ERP activity was computed as the average EEG amplitude across trials.

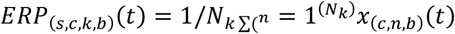

where *ERP*_(*s,c,k,b*)_(*t*) represents the event-related potential for subject *s*, channel *c*, task class *k*, and frequency band *b. N*_*k*_ is the number of trials for task class, and *x*_(*c,n,b*)_(*t*) is the filtered EEG signal from channel and trial. Separate ERP matrices were retained for all trials, left-hand motor imagery trials, and right-hand motor imagery trials.

The resulting ERP output for each subject and band had the form of a channel-by-time matrix.

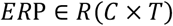

ERP features were used for descriptive visualization of cue-related temporal responses and were not treated as the primary inferential feature family for healthy-versus-stroke statistical comparison.

### Windowed Band Power

Windowed band-power features were extracted to compare baseline and motor imagery-period spectral activity across groups, channels, frequency bands, and task classes. Because the signals were already filtered into predefined frequency bands, this feature represented band-limited windowed power rather than broadband power spectral density. For each subject, channel, frequency band, task class, and analysis window, power was calculated as the mean squared amplitude of the bandpass-filtered signal within the selected time window.

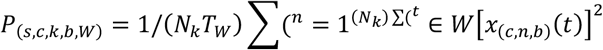

where *P*_(*s,c,k,b,W*)_ represents windowed band power for subject *s*, channel *c*, task class *k*, frequency band *b*, and analysis window *W*. The variable *T*_*W*_ represents the number of samples in the selected window, and *x*_(*c,n,b*)_(*t*) is the bandpass-filtered EEG signal.

For each band, power was retained separately for left-baseline, left-motor imagery, right-baseline, and right-motor imagery conditions.

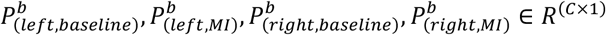

For the primary healthy-versus-stroke group comparison, left- and right-hand conditions were averaged within subject so that each participant contributed one baseline value and one motor imagery value per band and channel.

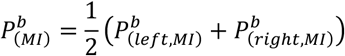

The resulting windowed band-power feature matrix had the form:

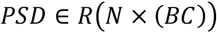

where *B* is the number of analyzed frequency bands and *C* is the number of channels. In this study, windowed band-power features were extracted for delta/slow-delta, theta, mu, beta, and gamma/high-frequency activity. Power values were retained in linear units for feature extraction and statistical analysis. For visualization, windowed band-power values were displayed as group means with standard-deviation error bars on a linear y-axis.

### Event-Related Desynchronization/Synchronization

Event-related desynchronization/synchronization was calculated to quantify task-related power modulation relative to baseline. ERD/ERS was emphasized in the mu and beta bands because these rhythms are closely associated with motor imagery and sensorimotor rhythm modulation.

ERD/ERS was computed using a trial-first approach. For each trial, instantaneous band-limited power was estimated by squaring the bandpass-filtered signal.

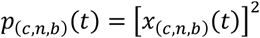

For each trial, baseline power was calculated as the mean power within the trial-specific baseline window.

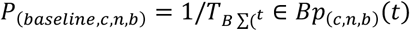

Motor imagery-window power was calculated as the mean power within the motor imagery window.

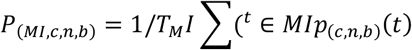

ERD/ERS was then calculated as percent change relative to baseline.

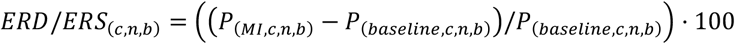

where *P*_*MI*_ represents mean power during the motor imagery window and *P*_*baseline*_ represents mean power during the baseline window for the same trial. Negative values indicate event-related desynchronization, whereas positive values indicate event-related synchronization.

For each subject, band, channel, and task class, trial-level ERD/ERS values were averaged to produce subject-level left-hand and right-hand motor imagery vectors.

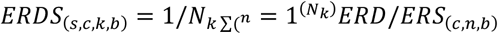

The resulting ERD/ERS feature matrix had the form:

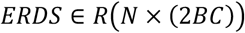

where *B* is the number of ERD/ERS frequency bands, *C* is the number of EEG channels, and the factor of 2 corresponds to left-hand and right-hand motor imagery conditions. In this study, ERD/ERS statistical features were extracted for the mu and beta bands. A moving-average smoothing window was applied only to plotting copies for visualization of ERD/ERS time-course heatmaps.

### Hemispheric Asymmetry Index

Hemispheric balance was evaluated using a normalized asymmetry index across homologous left-right electrode pairs. This feature was included because stroke can alter interhemispheric organization, lateralization, and recruitment of ipsilesional and contralesional motor networks during motor imagery.

For each subject, frequency band, task class, and electrode pair, asymmetry was calculated from the left- and right-hemisphere channel values.

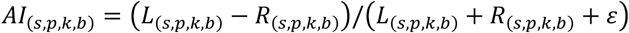

where *AI*_(*s,p,k,b*)_ represents the asymmetry index, *L*_(*s,p,k,b*)_ represents the left-hemisphere channel value, *R*_(*s,p,k,b*)_ represents the right-hemisphere channel value, and epsilon was added to prevent division by zero. Positive values indicate relatively greater left-hemisphere activity, whereas negative values indicate relatively greater right-hemisphere activity.

The following homologous electrode pairs were included: FP1-FP2, F3-F4, F7-F8, FC3-FC4, FT7-FT8, C3-C4, T3-T4, CP3-CP4, TP7-TP8, P3-P4, T5-T6, and O1-O2. Midline electrodes, including FZ, FCZ, CZ, PZ, and OZ, were not included because they do not have left-right homologous counterparts. Left-hand and right-hand motor imagery were retained separately because the expected direction and magnitude of lateralization may differ depending on the imagined hand.

The resulting ASSI feature matrix had the form:

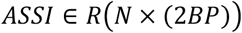

where *B* is the number of frequency bands, *P* is the number of homologous electrode pairs, and the factor of 2 corresponds to left-hand and right-hand motor imagery conditions.

### Time-Frequency Representations

Time-frequency representations were computed to characterize how motor imagery-related spectral activity changed across time. Unlike windowed band-power features, which reduce each trial to a single value within the baseline or motor imagery window, time-frequency analysis preserves both temporal and frequency information. This was important for the present study because acute stroke participants may show motor imagery-related responses that are delayed, transient, prolonged, or more variable than those observed in healthy participants.

Continuous wavelet transforms were used to estimate time-frequency activity. For each subject, time-frequency analysis was performed separately for left-hand and right-hand motor imagery trials and across the retained EEG channels. For a single channel and trial, the EEG signal was represented as: *X* {*c, n*}(*t*) where *c* represents channel, *n* represents trial, and *t* represents time. The continuous wavelet transform decomposes the signal by comparing it with scaled and shifted versions of a mother wavelet:

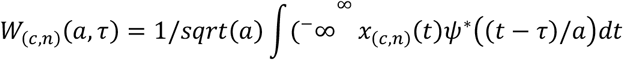

where (*W*_{*c,n*}_(*a, τ*)) is the wavelet coefficient, *a* is the wavelet scale, *τ* is time shift, and *ψ*^*^ is the complex conjugate of the mother wavelet. Morlet wavelets were used because they provide a practical balance between time and frequency localization for oscillatory EEG activity. In MATLAB, the continuous wavelet transform was implemented using the analytic Morlet wavelet option. Wavelet scale was converted to frequency so that the resulting coefficients could be represented as frequency-by-time matrices.

For each trial, wavelet power was calculated as the squared magnitude of the complex wavelet coefficients:

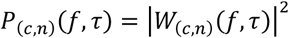

where *f* represents frequency and *τ* represents time. Power matrices were then averaged across trials within each motor imagery class:

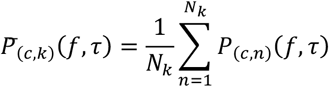

where *k* represents the task class, either left-hand or right-hand motor imagery, and *N*_*k*_ is the number of trials in that class.

To express task-related spectral activity relative to baseline, time-frequency power was baseline-normalized in decibels. For each frequency and channel, baseline power was calculated as the mean wavelet power within the baseline window:

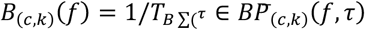

where *B* is the baseline window and *T*_*B*_ is the number of time samples in that window. Baseline-normalized time-frequency activity was then calculated as:

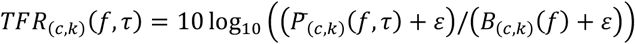

where *ϵ* was added to prevent division by zero. Positive values indicate increased power relative to baseline, whereas negative values indicate decreased power relative to baseline.

Time-frequency summaries were generated for the mu band, beta band, and broader 8–30 Hz sensorimotor range. For visualization, the baseline-normalized frequency-by-time matrix was averaged across the frequency bins within each range:

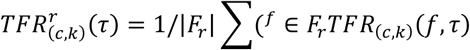

For healthy participant epochs containing NaN padding, the wavelet transform was computed only over the finite portion of the trial. If missing samples occurred within the finite segment, they were linearly interpolated before wavelet decomposition. The resulting wavelet power was then inserted back into the corresponding time locations of the full cue-aligned epoch. This preserved temporal alignment across subjects while preventing padded samples from influencing the wavelet transform.

Time-frequency representations were used as descriptive visualizations rather than as the primary statistical feature family. They were used to assess whether motor imagery-related spectral changes occurred during the expected task period and whether healthy and stroke participants differed in the timing, duration, or channel distribution of spectral modulation. A moving-average smoothing window was applied only to plotting copies of the channel-by-time summaries to improve figure readability; the underlying time-frequency values were not smoothed before computation.

### Group Averaging

Group-level summaries were generated from the saved subject-level feature files after feature extraction. Healthy and stroke participants were averaged separately for each feature family, including ERP, windowed band power, ERD/ERS, TFR, asymmetry index, and topographical/channel-power features. Group averages were used to visualize broad differences in motor imagery-related EEG patterns between neurologically intact and acute stroke participants.

For each feature, group averaging was performed across subjects rather than across pooled trials.

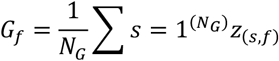

where *G*_*f*_ represents the group-average value for feature *f, N*_*G*_ is the number of subjects in group *G*, and *z*_(*s,f*)_ is the subject-level value for subject *s* and feature *f*. This ensured that each participant contributed equally to the group estimate.

Healthy group averages were used as the neurologically intact reference pattern. Stroke group averages were generated for comparison but were interpreted cautiously because acute stroke EEG may vary across participants due to lesion location, impairment severity, time after stroke, and individual neuroplastic reorganization. Therefore, group averages were used for descriptive visualization, while statistical testing was performed on subject-level feature values.

### Statistical Analysis

Statistical analysis was performed on subject-level EEG feature values so that each participant, rather than each trial, served as the statistical unit. Analyses were performed separately for each feature family, including windowed band power, ERD/ERS, asymmetry index, and topographical/channel-power features.

Within-group baseline versus motor imagery comparisons were performed separately for healthy and stroke participants using paired t-tests. These comparisons tested whether each group showed task-related feature changes during motor imagery relative to baseline. Between-group comparisons were performed using Welch’s two-sample t-tests because the healthy and stroke cohorts had unequal sample sizes and were not assumed to have equal variances (48). Welch’s tests were applied to both motor imagery-period values and motor imagery-minus-baseline responses, allowing group differences to be evaluated for absolute motor imagery activity and task-related modulation.

Benjamini-Hochberg false discovery rate correction was applied within each feature family and comparison type to account for multiple comparisons across channels, frequency bands, task classes, and feature types (49). Effect sizes were calculated to support interpretation of the magnitude of group-level differences in addition to statistical significance (50).

### Window-Size Comparison and Transferability Ranking

After subject-level feature extraction and healthy-versus-stroke statistical testing were completed separately for the 0.5 s and 1.0 s analysis windows, a post hoc window-comparison analysis was performed to evaluate which EEG features were most suitable for healthy-to-stroke transfer learning. This analysis was designed to address whether a longer post-onset feature window improved transfer-relevant feature stability in the stroke population, rather than simply identifying features that differed significantly between groups.

For each feature family included in the inferential analysis, including PSD, ERD/ERS, ASSI, and TOPO/channel-power features, the group-average statistics tables from the 0.5 s and 1.0 s windows were loaded and matched by feature name. Each feature therefore had paired summary values across the two analysis windows, including healthy MI mean, stroke MI mean, healthy MI standard error, stroke MI standard error, healthy-minus-stroke MI difference, and FDR-based significance status from the corresponding healthy-versus-stroke MI comparison.

Feature transferability was evaluated using three related criteria: healthy-stroke similarity, window stability, and group variability. Healthy-stroke similarity was quantified by the normalized absolute difference between the healthy MI mean and stroke MI mean, scaled by the average magnitude of the two group means. A smaller normalized gap indicated greater similarity between populations. Window stability was quantified by measuring how much the healthy and stroke group means changed between the 0.5 s and 1.0 s windows. Group variability was estimated using the ratio between the standard error of the mean and the corresponding group mean magnitude. Features with smaller healthy-stroke differences, smaller window-dependent shifts, and lower relative variability were considered stronger candidates for healthy-to-stroke transfer.

A transferability score was computed separately for the 0.5 s and 1.0 s windows and then averaged to produce an overall transferability score. Higher transferability scores indicated features that were more similar between healthy and stroke participants and less variable across groups. A separate longer-window benefit score was calculated to determine whether the 1.0 s window improved transfer suitability relative to the 0.5 s window. This score increased when the 1.0 s window improved healthy-stroke similarity, reduced stroke variability, or increased the feature’s transferability score relative to the shorter window. Therefore, the longer-window analysis directly tested whether stroke participants may require a longer temporal window for stable MI feature extraction.

A discriminative score was also computed to identify features that most strongly separated healthy and stroke groups. This score emphasized healthy-stroke feature separation and incorporated the FDR-based significance status from the healthy-versus-stroke MI comparison. Discriminative features were interpreted separately from transfer-suitable features. Features with high discriminative scores were considered useful for characterizing stroke-related EEG differences, whereas features with high transferability scores were considered more appropriate candidates for healthy-to-stroke transfer initialization. This distinction was important because a feature that strongly differentiates healthy and stroke participants may reveal clinically relevant stroke-related changes, but may not be optimal for direct transfer without target-specific adaptation.

For each feature family, features were ranked separately by overall transferability, longer-window benefit, and healthy-versus-stroke discriminability. Combined rankings across all feature families were also generated. These rankings were used to identify EEG features that may provide a shared neural representation for initializing an adaptive decoder in acute stroke participants, as well as features that may require stroke-specific adaptation.

### BMI/Q-KTD Time-Series Input Preparation

In addition to generating group-level features, motor imagery epochs were generated as trial-level time-series inputs. For each subject, motor imagery band epochs were segmented relative to motor imagery onset and saved as both a three-dimensional input tensor and a flattened input matrix. The tensor preserved the trial-by-time-by-channel structure, while the flattened matrix provided a trial-by-feature representation for downstream decoder training and Q-KTD analysis.

A 0–0.5 s post-onset motor imagery window was generated for each participant and used as the Q-KTD input. The 0.5 s window was used as the primary early-response input window because it captures the immediate post-cue EEG activity while limiting the amount of post-onset data included in the decoder input.

Each BMI/Q-KTD file retained the subject identifier, group label, sampling rate, channel labels, trial labels, selected time window, input tensor, and flattened input matrix. Separate files were saved for each time window using window-specific filenames so that the 0.5 s inputs could be compared directly without overwriting or mixing decoder inputs.

### Decoder Feature Organization

For decoder analysis, trial-level EEG features were organized so that each trial corresponded to a left-hand or right-hand motor imagery class. Feature matrices retained participant identifiers, group labels, trial numbers, and class labels. This structure allowed the same processed dataset to support both physiological comparison and Q-KTD decoding.

The physiological analyses were used to characterize shared and altered EEG structure between healthy and stroke participants, while the decoder features were used to test whether source-population EEG information improved adaptive classification of left-versus right-hand motor imagery in stroke participants.

### Q-KTD

For Q-KTD decoding, trial-level EEG inputs were structured around the beginning of the MI period to approximate early online BMI behavior. Each decoding trial used the cue-aligned epoch, but the decoder input feature was extracted from an early 0.5 s post-cue MI segment beginning at cue onset.

A reinforcement-learning decoder was used because assistive BMI control requires adaptation to noisy, nonstationary, and user-specific neural signals. In this framework, EEG features were treated as states, decoder outputs as actions, and task feedback as reward. This structure allowed decoder learning to be evaluated as an adaptive control problem rather than only as static classification (45–50).

In the reinforcement-learning BMI framework, the decoder functions as an agent that works with the user to achieve a goal. Neural signals at time *t* are represented as states *x* ∈ *X*, and the decoder selects an action *a ϵ A*, where *X* and *A* represent the possible state and action spaces. The selected action changes the environment and generates a new neural state at time *t* + 1. Based on the state of the BMI device relative to the task goal, a reward, *r*(*t* + 1), is assigned to the agent. This process repeats until the task reaches a terminal state or the trial is completed (48).

Because EEG feature vectors are continuous and high-dimensional, the Q-function cannot be represented efficiently using a simple tabular form. Kernel Temporal Difference learning was therefore used to approximate the action-value function with nonlinear function approximation. The Q-function was represented using kernel expansion:

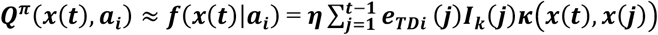

Where *K*(*x*(*t*), *x*(*j*)) is the kernel similarity between the current EEG state *x*(*t*) and a stored kernel center *x*(*j*)_*TD,j*_ represents the learned temporal difference weight, and *I*_*k*_(*j*) indicates the action associated with the *j*-th kernel unit (51, 52).

The temporal-difference error was used to update the action-value estimate:

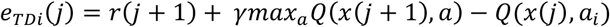

Where *r*(*j* + 1) is the reward received after action selection, and the second term estimates the discounted future value of the next state. This update allowed the decoder to adjust its action-value function based on prediction error between expected and observed reward (38,39).

A Gaussian kernel was used for nonlinear feature mapping. Because kernel methods can grow in complexity as new samples are added, quantization was used to limit kernel expansion. A new kernel center was added only when the minimum distance between the current EEG input and all existing kernel centers exceeded a predefined threshold:

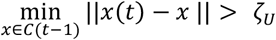

Where *C*(*T* − 1) is the set of existing kernel centers and *ζ*_*U*_is the quantization threshold (39,42).

Feature vectors were normalized to the range of −1 to 1 before decoding because EEG features differed in scale across subjects and datasets. Actions were selected using an epsilon-greedy policy because reinforcement learning requires a balance between exploiting learned action values and exploring alternative actions. The decoder selected the action with the highest estimated Q-value with probability 1-*ϵ*, while selecting a random exploratory action with probability *ϵ*. The step size was set to *η* = 0.5, the discount factor was set to *γ* = 0.9, and the exploration rate was set to *ϵ* = 0.01. The quantization threshold was set to *ζ*_*U*_ = 0.1 for all cases.

### Transfer-Learning

Transfer learning was evaluated because the main assistive technology question was whether prior information from a source population could reduce early decoder burden for stroke users. Healthy-to-stroke transfer tested whether healthy MI EEG contained useful structure for initializing stroke-subject decoding.

For healthy-to-stroke transfer, the Q-KTD model was first trained on the combined healthy MI EEG data until convergence. Learned kernel centers and associated model weights from the healthy source model were then transferred to initialize each stroke-subject target model (34,43). Each stroke participant was evaluated under two conditions: Q-KTD without transfer learning and Q-KTD with healthy-to-stroke transfer learning.

### Experimental Setting

Offline datasets do not provide true human-in-the-loop BMI interaction, so decoding was evaluated using a simulated center-out reaching task. This task approximated a practical BMI setting in which neural decoding outputs are mapped to movement of an external device. At the start of each trial, the agent’s cursor began at the center of a circle with radius 1. Class targets were placed along the circumference, with one target designated as the goal for each trial. The target location corresponded to the MI class for that trial.

A reward was assigned based on the Euclidean distance between the updated cursor location and the goal. If the cursor reached within 0.1 of the goal, the agent received a positive reward equal to the total number of targets minus 1. If the cursor did not reach the goal, the agent received a reward of −1. This simulated the relationship between BMI classification output and external device movement used in practical BMI control tasks.

Although the setup did not allow real user co-adaptation because prerecorded datasets were used, the algorithm was applied in an online-style manner. Each trial corresponded to a single decision step in which the agent received an EEG feature vector, selected an action, and received a reward based on the resulting cursor position.

Performance EvaluationThe Q-KTD algorithm was run for 10 Monte Carlo repetitions for each decoding condition because random trial ordering and random model initialization can influence reinforcement-learning performance. Each repetition used randomized trial ordering and randomized initial model weights. Models were allowed to run for up to 50 epochs. A stopping criterion was applied such that learning stopped when the average success rate across the current epoch and two previous epochs did not improve by more than 0.01.

Success rate was calculated as the number of successful trials divided by the total number of trials. First-epoch success rate was used as the primary performance measure because early decoder performance is most relevant for practical BMI use, where the system must become useful quickly and extensive subject-specific calibration may limit adoption. Later-epoch performance was also examined to determine whether transfer learning primarily improved early adaptation or final convergence.

### Statistical Analysis

For the transfer-learning (TL) analysis, first-epoch success rates were compared across Monte Carlo repetitions within each stroke participant. Paired t-tests (p<0.01) were run between TL versus non-TL results for stroke individuals (with healthy population models as the source domain).

## Results

### Motor Imagery Event-related Potentials

Figure 1 presents representative C3 event-related potentials from one healthy participant (S004) and one acute stroke participant (sub-04) during left- and right-hand motor imagery. In the healthy example, motor imagery onset occurred at 0 s, with the baseline defined immediately before onset and the 0–0.5 s interval used as the motor imagery analysis window. In the stroke example, motor imagery began at 2 s, while the baseline was taken from the late-trial rest period, consistent with the dataset-specific experimental paradigm.

**Figure 1.**
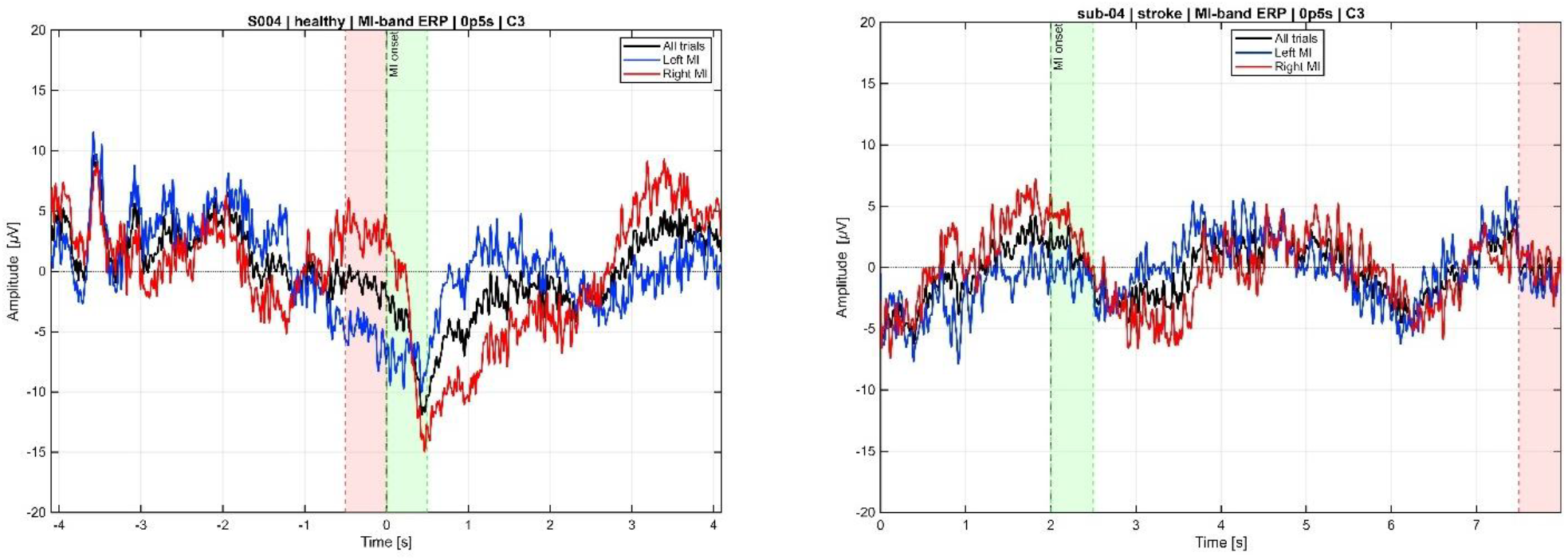
Representative C3 event-related potentials during motor imagery in healthy and acute stroke participants. Left: representative healthy participant S004. Right: representative acute stroke participant sub-04. Trial-averaged waveforms are shown for all motor imagery trials, left-hand motor imagery, and right-hand motor imagery. Shaded regions indicate the dataset-specific baseline and 0.5 s motor imagery analysis windows. Motor imagery onset occurred at 0 s for the healthy dataset and at 2 s for the stroke dataset, reflecting differences in the original experimental paradigms.

As shown in Figure 1A, the healthy participant exhibited a pronounced negative deflection after motor imagery onset, with visible separation between the left- and right-hand motor imagery waveforms during the early post-onset interval. The right-hand motor imagery response showed the larger negative deflection within the 0.5 s window. In contrast, the stroke participant in Figure 1B showed a smaller and more gradual post-onset change at C3. Left- and right-hand motor imagery responses were less clearly separated within the first 0.5 s, with greater divergence appearing later in the motor imagery period.

These representative examples illustrate that motor imagery-related temporal modulation was present in both datasets, while the magnitude, timing, and class-specific waveform patterns differed between participants. Because the plots represent individual participants, they were used for descriptive visualization rather than group-level statistical inference.

### Baseline and post-cue spectral power across EEG channels

Figure 2 shows group-averaged mu-band power during the dataset-specific baseline and motor imagery periods using the 0.5 s analysis window. Healthy and acute stroke participants showed broadly overlapping mu-band power values across most channels. The stroke group exhibited greater interparticipant variability at several locations, with a particularly large standard deviation at CP3 during motor imagery. However, no channel-level healthy–stroke differences in mu-band power remained statistically significant after false discovery rate correction.

**Figure 2.**
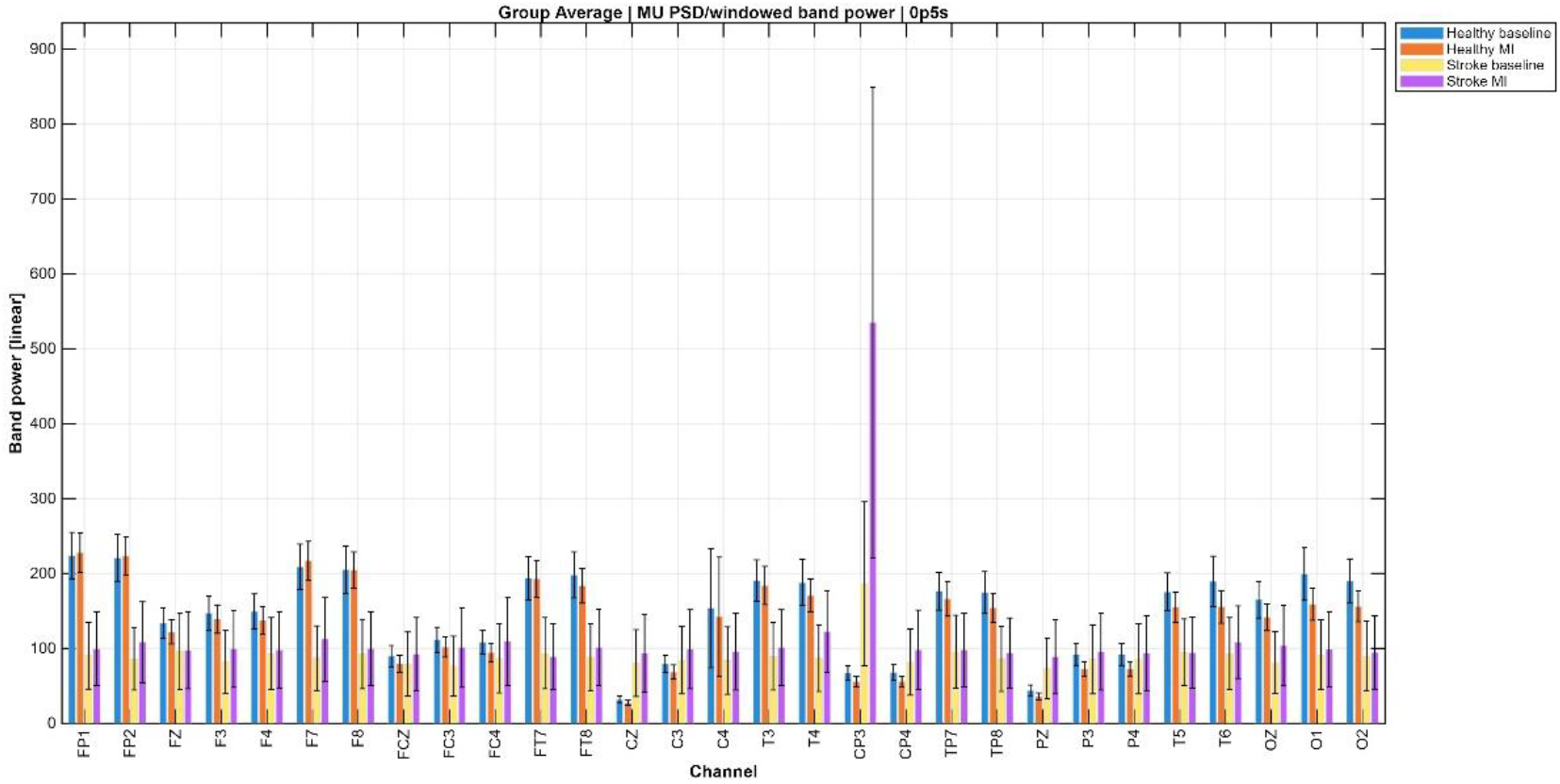
Group-averaged mu-band power in healthy and acute stroke participants. Windowed mu-band power was calculated over the 0.5 s dataset-specific baseline and motor imagery periods across the harmonized EEG montage. For each channel, bars represent healthy baseline, healthy motor imagery, stroke baseline, and stroke motor imagery values. Error bars represent standard deviations across participants. No channel-level healthy–stroke differences remained statistically significant after false discovery rate correction.

The additional frequency-band analyses showed similar overlap between the healthy and stroke groups. Delta/slow-delta power tended to be higher in the stroke group during motor imagery across several channels, while beta-band power also showed heterogeneous baseline and motor imagery values in the stroke cohort. Theta- and gamma-band results followed comparable patterns of substantial group overlap and greater variability among stroke participants. Nevertheless, no channel-level healthy–stroke differences survived multiple-comparison correction in any of the evaluated frequency bands.

The 1.0 s analyses produced broadly similar results to the 0.5 s analyses. Although the longer window changed the magnitude of selected channel-level means and standard deviations, it did not reveal statistically significant healthy– stroke differences after correction. Overall, the windowed band-power results indicate considerable overlap in group-level power distributions, together with greater heterogeneity in the acute stroke cohort.

### ERD/ERS during motor imagery

Figure 3 compares ERD/ERS values between healthy and acute stroke participants during left- and right-hand motor imagery using the 0.5 s analysis window. Positive ERD/ERS values predominated across most features in both groups, indicating that early task-related power enhancement was more common than power suppression under the applied sign convention.

**Figure 3.**
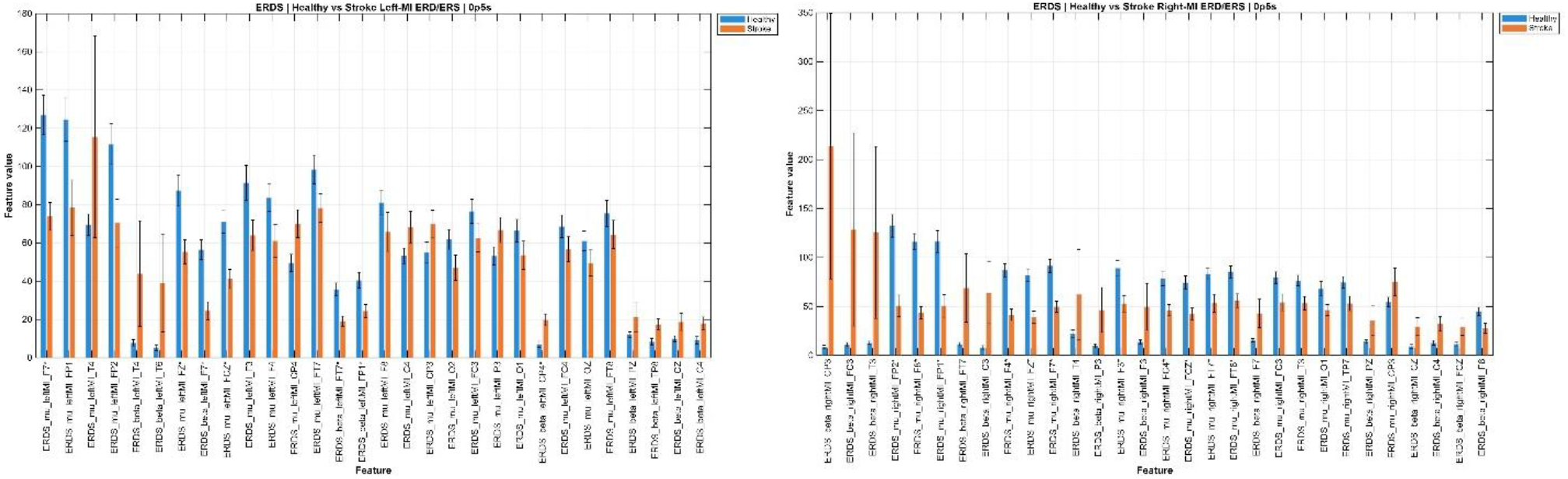
Group-level ERD/ERS values in healthy and acute stroke participants during motor imagery. Left: ERD/ERS values during left-hand motor imagery using the 0.5 s analysis window. Right: ERD/ERS values during right-hand motor imagery using the 0.5 s analysis window. Bars represent group means for healthy and acute stroke participants, and error bars represent standard deviations across participants. Negative values indicate event-related desynchronization, whereas positive values indicate event-related synchronization. The y-axis scales differ between the left- and right-hand motor imagery panels to accommodate differences in the observed value ranges.

For both left- and right-hand motor imagery, healthy participants generally showed larger ERD/ERS values across several frontal, frontocentral, central, and centroparietal features. The stroke group showed a more heterogeneous pattern, with lower values across many features but larger responses at selected channels. Standard deviations were frequently larger in the stroke group, indicating greater interparticipant variability in task-related spectral modulation. Although the overall healthy–stroke relationship was similar for left- and right-hand motor imagery, the magnitude and spatial distribution of the responses differed between the two imagined-hand conditions.

The 1.0 s analysis showed a less consistent group relationship and greater variability than the 0.5 s analysis. Several stroke-group features exhibited markedly larger positive ERD/ERS values and wider standard deviations when the longer window was used. For left-hand motor imagery, these larger values were most apparent around central and centroparietal features, including CP3, CP4, C4, Cz, T4, and Pz. For right-hand motor imagery, prominent increases were observed particularly around CP3 and Pz. The longer analysis window therefore captured additional task-related activity but also produced more extreme and variable ERD/ERS estimates in the stroke group. Overall, the comparison indicates that ERD/ERS magnitude and variability were sensitive to the selected analysis-window duration.

### Time-frequency organization of motor imagery activity

Figures 4 and 5 show channel-by-time summaries of baseline-normalized mu-band time–frequency power for healthy and acute stroke participants, respectively. Negative values indicate reduced mu power relative to baseline, whereas positive values indicate increased mu power.

**Figure 4.**
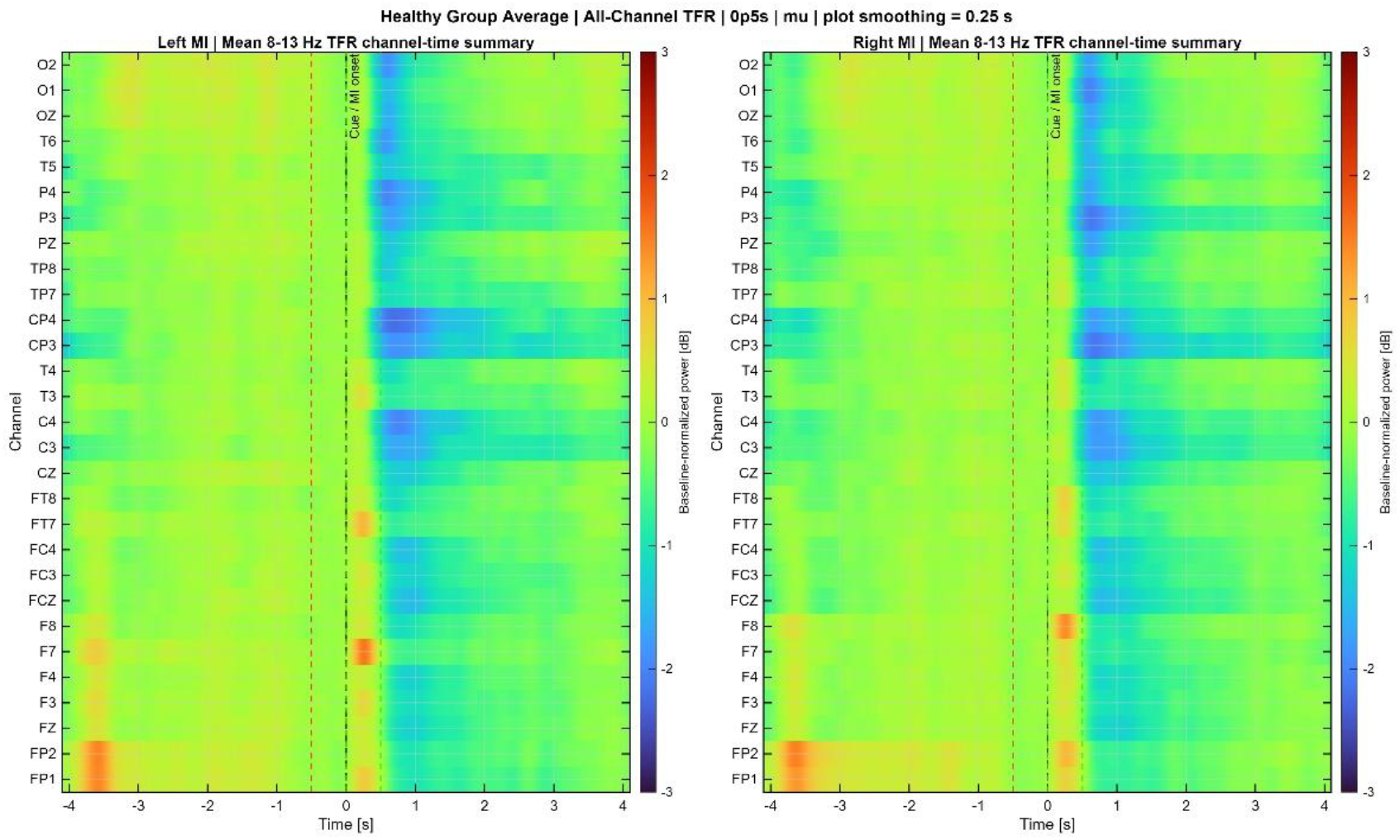
Group-averaged mu-band time–frequency activity during motor imagery in healthy participants. Left: baseline-normalized mu-band activity during left-hand motor imagery. Right: baseline-normalized mu-band activity during right-hand motor imagery. Rows represent the harmonized EEG channels, and color represents baseline-normalized power in decibels using the 0.5 s analysis window. Motor imagery onset occurred at 0 s. Negative values indicate reduced mu-band power relative to baseline, whereas positive values indicate increased power. Vertical reference lines indicate the dataset-specific motor imagery onset and analysis periods.

**Figure 5.**
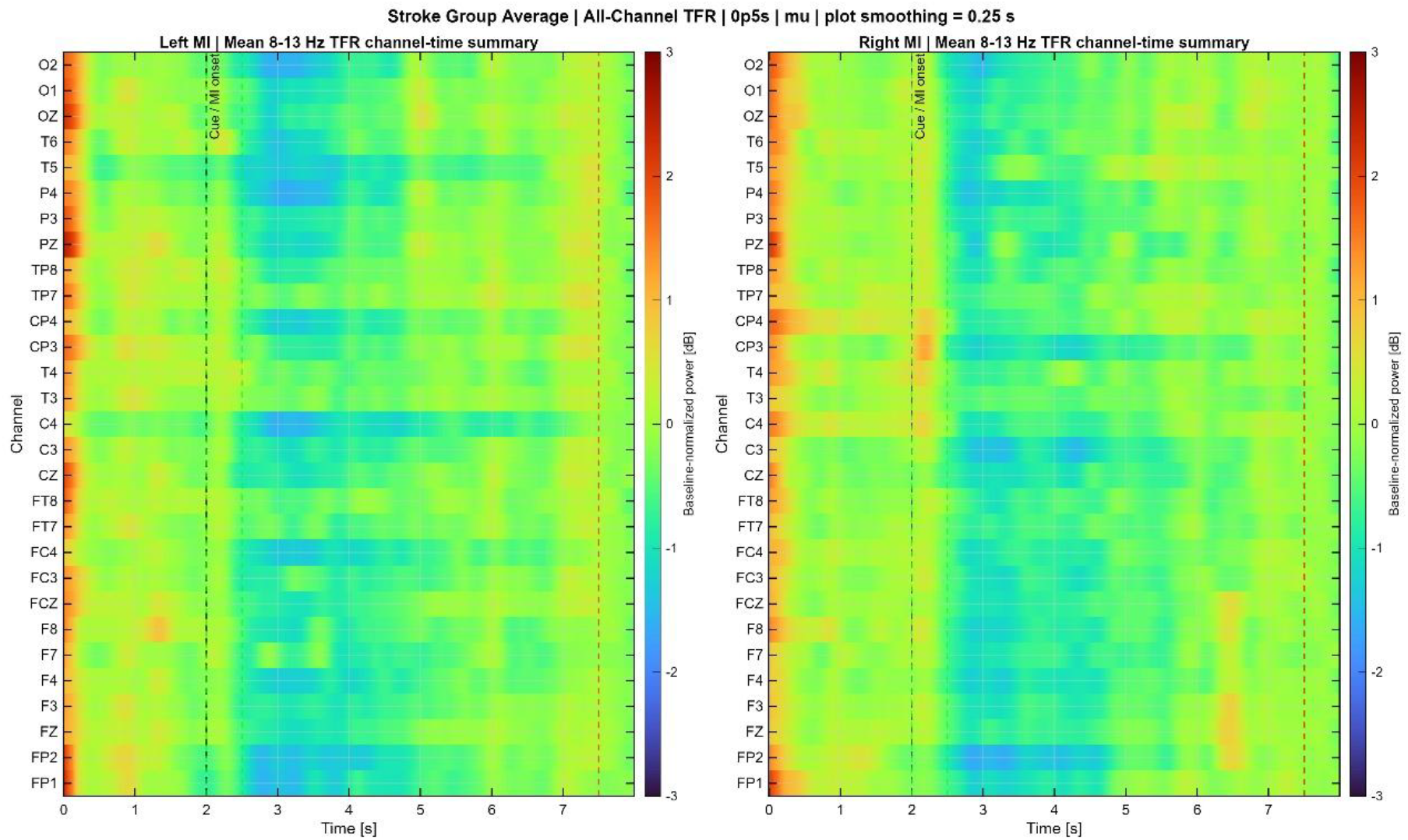
Group-averaged mu-band time–frequency activity during motor imagery in acute stroke participants. Left: baseline-normalized mu-band activity during left-hand motor imagery. Right: baseline-normalized mu-band activity during right-hand motor imagery. Rows represent the harmonized EEG channels, and color represents baseline-normalized power in decibels using the 0.5 s analysis window. Motor imagery onset occurred at 2 s according to the original stroke-task paradigm. Negative values indicate reduced mu-band power relative to baseline, whereas positive values indicate increased power. Vertical reference lines indicate the dataset-specific motor imagery onset, analysis period, and late-trial baseline period.

In the healthy group, shown in Figure 4, both left- and right-hand motor imagery produced a clear reduction in mu-band power following motor imagery onset at 0 s. The suppression emerged shortly after onset and was most pronounced during approximately the first 1–2 s of the motor imagery period. The strongest negative modulation was observed across central and centroparietal channels, including C3, C4, CP3, and CP4, with additional suppression extending into frontal and posterior channels. Left- and right-hand motor imagery showed broadly similar temporal patterns, although the magnitude and channel distribution differed between conditions. The group-level responses did not display a simple mirror-symmetric lateralization pattern.

The acute stroke group also exhibited post-onset mu-band suppression during both left- and right-hand motor imagery, as shown in Figure 5. Following motor imagery onset at 2 s, negative modulation developed across central, frontocentral, centroparietal, and posterior channels. Compared with the healthy-group pattern, the stroke response appeared more spatially diffuse and temporally heterogeneous. Suppression varied across channels in its onset, magnitude, and duration, and localized periods of increased power were interspersed with the negative modulation. The stroke-group response also appeared to persist across a broader portion of the motor imagery interval, with a less uniform return toward baseline.

The corresponding 1.0 s analyses preserved the principal patterns observed with the 0.5 s window. In the healthy group, post-onset mu suppression remained organized across central and centroparietal regions for both imagined-hand conditions. In the stroke group, the longer window continued to show broader and more variable channel-level modulation. The 1.0 s window altered the magnitude and contrast of selected responses but did not reveal a fundamentally different temporal or spatial organization. Thus, the major mu-band patterns were qualitatively stable across the two analysis-window durations.

### Hemispheric asymmetry during motor imagery

Figure 6 compares hemispheric asymmetry between healthy and acute stroke participants during the 0.5 s motor imagery window. The displayed features include multiple frequency bands, imagined-hand conditions, and homologous left–right electrode pairs. Positive values indicate relatively greater left-hemisphere activity, whereas negative values indicate relatively greater right-hemisphere activity.

**Figure 6.**
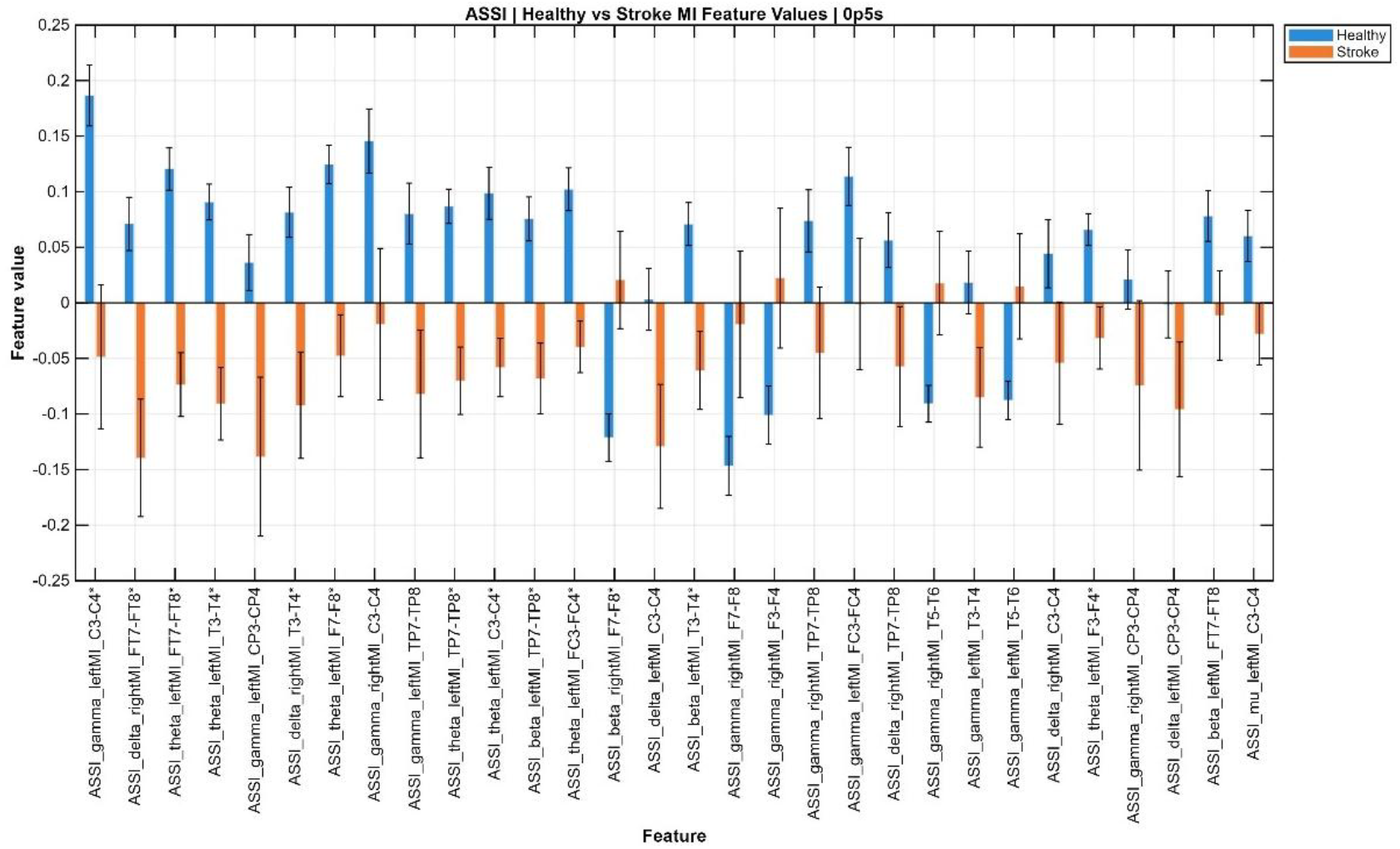
Hemispheric asymmetry during motor imagery in healthy and acute stroke participants. Asymmetry-index values are shown for selected frequency bands, imagined-hand conditions, and homologous left–right electrode pairs using the 0.5 s motor imagery window. Bars represent group means for healthy and acute stroke participants, and error bars represent standard deviations across participants. Positive values indicate relatively greater left-hemisphere activity, whereas negative values indicate relatively greater right-hemisphere activity. Asterisks indicate statistically significant healthy–stroke differences after false discovery rate correction.

Across many features, healthy participants showed positive asymmetry values while stroke participants showed negative values. This pattern was observed across several frontal, frontotemporal, central, and centroparietal electrode pairs and was present in multiple frequency bands during both left- and right-hand motor imagery. Several features therefore showed an apparent reversal in the direction of hemispheric balance between the two groups.

The group relationship was not uniform across all features. Some electrode-pair and frequency-band combinations showed negative asymmetry in healthy participants and values closer to zero or positive in the stroke group. The findings therefore indicate feature-dependent alteration of hemispheric organization after stroke rather than a consistent shift toward one hemisphere. Greater variability was also observed in the stroke group for several features, consistent with heterogeneous post-stroke neural organization.

The corresponding 1.0 s motor imagery analysis showed broadly similar directions of asymmetry for many features, although the magnitude of selected values changed with the longer window. In contrast, the 1.0 s motor imagery-minus-baseline analysis showed greater group overlap and more values near zero. Overall, the main healthy–stroke differences in absolute motor imagery asymmetry were qualitatively preserved across the two window durations.

### Healthy-to-stroke Q-KTD transfer-learning performance

For both cases, with and without TL implemented, Q-KTD was able to map EEG motor imagery features to movement intention during the first training epoch, though moderate variability was observed between subjects. Across the 50 stroke subjects, TL produced a modest overall improvement in first-epoch success rate. The average first-epoch success rate increased from 49.46% without TL to 51.82% with TL.

However, the effect of TL was not uniform across the stroke cohort. The observed subject-level changes ranged from −14.0% to 18.75%, as listed in Table 1. A total of 29 subjects showed improved first-epoch performance with TL, with an average increase of 7.3%, while 21 subjects showed decreased performance. The largest improvements were observed for sub-14, sub-15, sub-48, sub-26, and sub-23, with increases of 18.75%, 17.0%, 15.75%, 15.25%, and 14.25%, respectively. These results suggest that, for some subjects, transferring learned information from the healthy cohort provided a strong initialization that improved early-stage learning.

**Table 1.** Subject-level first-epoch Q-KTD success rates with and without healthy-source transfer learning. Asterisks indicate p<0.01 based on paired tests across the 10 Monte Carlo runs.

| Subject | Without TL (%) | With TL (%) | Change (%) |
| --- | --- | --- | --- |
| sub-01 | 59.50 | 54.00 | -5.50 |
| sub-02 | 46.75 | 45.75 | -1.00 |
| sub-03 | 59.00 | 46.75 | -12.25* |
| sub-04 | 48.50 | 50.75 | 2.25 |
| sub-05 | 48.50 | 54.75 | 6.25 |
| sub-06 | 50.75 | 47.00 | -3.75 |
| sub-07 | 49.25 | 50.00 | 0.75 |
| sub-08 | 52.50 | 47.50 | -5.00 |
| sub-09 | 60.50 | 58.50 | -2.00 |
| sub-10 | 44.75 | 55.50 | 10.75* |
| sub-11 | 49.00 | 48.50 | -0.50 |
| sub-12 | 57.75 | 52.00 | -5.75* |
| sub-13 | 47.75 | 55.00 | 7.25 |
| sub-14 | 50.25 | 69.00 | 18.75* |
| sub-15 | 42.00 | 59.00 | 17.00* |
| sub-16 | 43.75 | 44.25 | 0.50 |
| sub-17 | 49.75 | 49.00 | -0.75 |
| sub-18 | 46.00 | 53.75 | 7.75 |
| sub-19 | 51.00 | 46.50 | -4.50 |
| sub-20 | 45.50 | 46.75 | 1.25 |
| sub-21 | 54.25 | 53.75 | -0.50 |
| sub-22 | 53.25 | 60.25 | 7.00* |
| sub-23 | 45.75 | 60.00 | 14.25* |
| sub-24 | 42.00 | 48.25 | 6.25 |
| sub-25 | 55.50 | 62.50 | 7.00* |
| sub-26 | 43.00 | 58.25 | 15.25* |
| sub-27 | 47.50 | 37.75 | -9.75* |
| sub-28 | 56.25 | 56.00 | -0.25 |
| sub-29 | 55.50 | 54.50 | -1.00 |
| sub-30 | 68.75 | 58.25 | -10.50* |
| sub-31 | 49.25 | 49.00 | -0.25 |
| sub-32 | 50.00 | 45.00 | -5.00 |
| sub-33 | 47.75 | 53.00 | 5.25 |
| sub-34 | 35.75 | 44.25 | 8.50* |
| sub-35 | 57.00 | 60.00 | 3.00 |
| sub-36 | 44.25 | 54.25 | 10.00* |
| sub-37 | 40.00 | 43.50 | 3.50 |
| sub-38 | 54.00 | 50.50 | -3.50 |
| sub-39 | 40.75 | 47.25 | 6.50 |
| sub-40 | 52.50 | 56.50 | 4.00 |
| sub-41 | 62.00 | 48.00 | -14.00* |
| sub-42 | 46.75 | 41.75 | -5.00 |
| sub-43 | 44.25 | 45.50 | 1.25 |
| sub-44 | 55.75 | 58.75 | 3.00 |
| sub-45 | 49.75 | 59.50 | 9.75* |
| sub-46 | 38.50 | 45.75 | 7.25* |
| sub-47 | 37.25 | 48.00 | 10.75* |
| sub-48 | 40.25 | 56.00 | 15.75* |
| sub-49 | 50.50 | 52.75 | 2.25 |
| sub-50 | 52.00 | 47.75 | -4.25 |

In contrast, several subjects showed reduced first-epoch performance when TL was applied. The largest decreases were observed for sub-41, sub-03, sub-30, sub-27, and sub-12, with changes of −14.0%, −12.25%, −10.5%, −9.75%, and −5.75%, respectively. This indicates that although healthy-to-stroke TL can improve early decoding performance for many subjects, it may also negatively affect performance when the transferred source information does not align well with the target subject’s EEG representation.

In the offline analysis, the neural decoder encounters the same dataset across epochs; therefore, performance at the first epoch reflects the expected initial behavior of the decoder before repeated exposure to the same data. As such, the first epoch serves as a proxy for early closed-loop BMI performance, where rapid adaptation is critical. The updated results show that TL can improve this early learning stage for a majority of subjects, but the variability across subjects suggests that TL should be evaluated on an individual basis rather than assumed to be universally beneficial.

Figure 7 presents the average first-epoch success rates for all 50 stroke subjects, comparing TL and non-TL conditions. Error bars represent the standard deviation across the 10 Monte Carlo runs, and asterisks indicate subjects with statistically significant differences between TL and non-TL performance. The figure highlights the subject-specific nature of the TL effect. Several subjects, including sub-14, sub-15, sub-23, sub-26, sub-45, sub-47, and sub-48, showed clear improvement with TL, while other subjects, including sub-03, sub-12, sub-27, sub-30, and sub-41, showed significant decreases. Overall, the figure supports the numerical trend shown in Table 1: TL improved first-epoch performance for more than half of the stroke subjects, but negative transfer occurred in a meaningful subset of cases.

**Figure 7.**
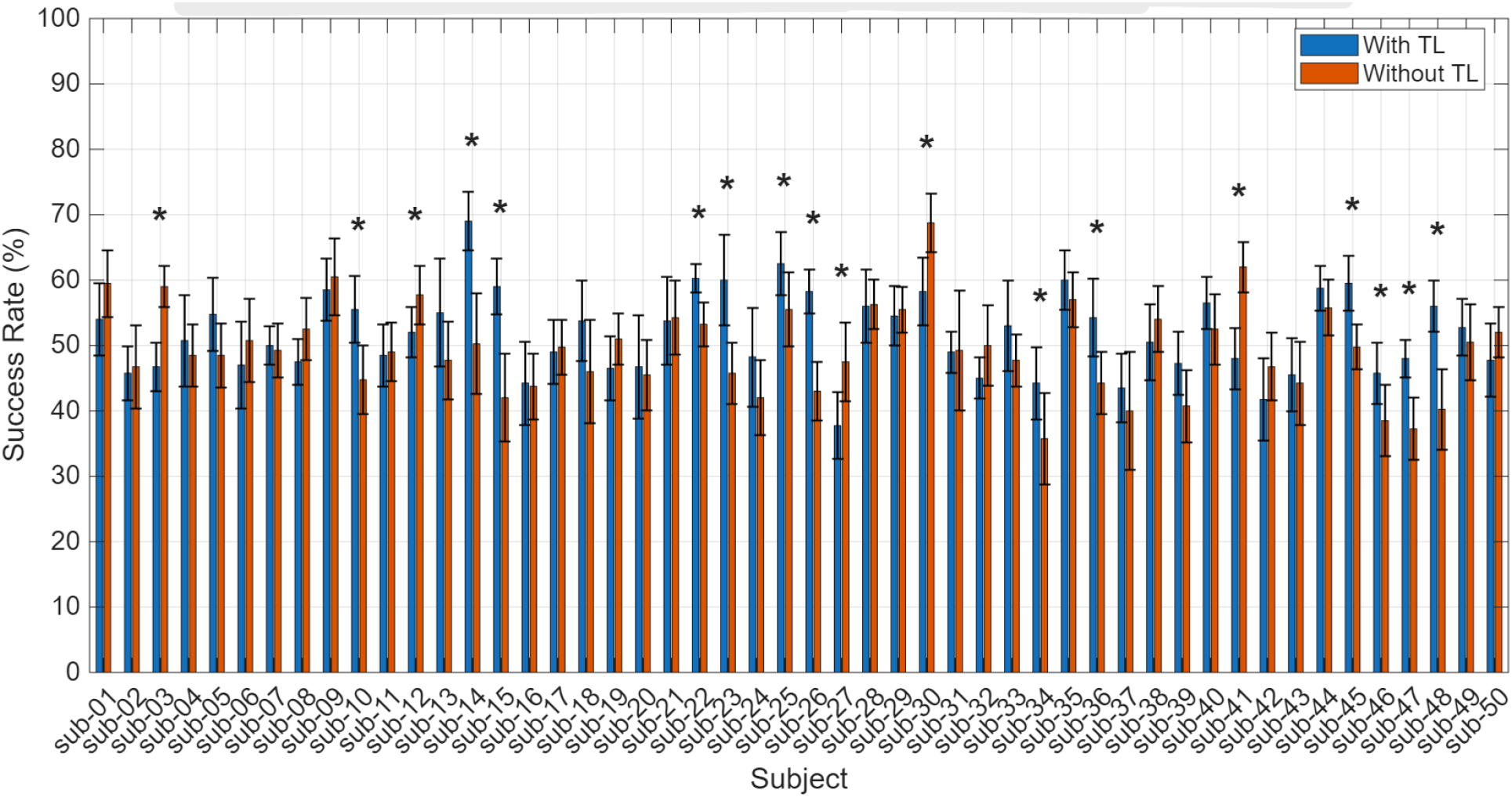
Subject-level first-epoch Q-KTD success rates with and without healthy-source transfer learning. Mean first-epoch motor imagery decoding success rates are shown for each of the 50 acute stroke participants across 10 Monte Carlo runs. Blue bars represent decoding with healthy-source transfer learning, and orange bars represent decoding without transfer learning. Error bars indicate standard deviations across Monte Carlo runs. Asterisks indicate statistically significant differences between the transfer-learning and no-transfer conditions based on participant-level paired t-tests (p<0.01).

## Discussion

This study examined whether motor imagery EEG learned from a healthy source population could support early Q-KTD decoding in individuals with acute stroke. The principal findings were that healthy and stroke participants retained some shared motor imagery–related EEG organization, while the stroke cohort showed greater interparticipant variability and altered temporal and hemispheric patterns. Healthy-source transfer learning increased mean first-epoch Q-KTD success rate from 49.46% to 51.82%, corresponding to an average improvement of 2.36%. Positive transfer occurred in 29 of 50 participants, whereas 21 participants showed reduced performance following transfer. These findings indicate that healthy-population EEG can provide useful initialization for some stroke participants, but that the benefit is strongly dependent on the individual target domain.

The physiological analyses provide context for this variable transfer response. Windowed band-power distributions overlapped substantially between the healthy and stroke groups, and no channel-level band-power differences remained significant after false discovery rate correction. This overlap suggests that the stroke data retained some frequency-specific structure observed in the healthy population. At the same time, the stroke group generally showed larger standard deviations and more heterogeneous baseline and motor imagery power across channels. Therefore, the absence of corrected group-level differences should not be interpreted as equivalence between the populations. Rather, the results indicate that broad group means may overlap while individual stroke participants express substantially different EEG patterns.

The time–frequency results provided additional evidence of both shared and altered motor imagery activity. Healthy and stroke participants showed post-onset mu-band power reductions during left- and right-hand motor imagery, indicating preservation of task-related sensorimotor modulation in both populations. In healthy participants, this modulation appeared relatively organized across central and centroparietal channels. In the stroke group, the response was more spatially diffuse and varied more across channels in its onset, magnitude, and duration. The corresponding 1.0 s analyses produced qualitatively similar patterns, suggesting that the principal temporal organization was not restricted to the 0.5 s window. However, because the two window durations were compared descriptively rather than through a formal statistical test, these observations should be interpreted as qualitative stability rather than evidence of window-size invariance.

The ERD/ERS and hemispheric-asymmetry findings further demonstrated that post-stroke motor imagery activity was not characterized by a uniform loss of sensorimotor modulation. Positive ERD/ERS values predominated during the early analysis windows in both groups, although stroke responses were more variable and showed larger values at selected channels, particularly in the 1.0 s analysis. Hemispheric asymmetry also differed in both magnitude and direction across several frequency-band and electrode-pair combinations. Some features showed positive asymmetry in the healthy group and negative asymmetry in the stroke group, whereas other features showed the opposite relationship. These results are more consistent with feature-dependent reorganization of hemispheric balance than with a single generalized shift toward either hemisphere. Because lesion laterality was not incorporated into the present analysis, the observed left–right asymmetry should not be interpreted directly as ipsilesional or contralesional recruitment.

The Q-KTD findings extend previous work demonstrating that kernel temporal-difference learning can acquire mappings between EEG motor imagery states and BMI control actions through repeated reward-based learning (38,39). Earlier EEG-based Q-KTD work showed that performance can improve substantially over repeated epochs, but also identified slow early learning as an important barrier to practical implementation (39). Transfer learning addresses this limitation by providing an inductive initialization rather than requiring the target model to begin with minimal prior information. In the present study, kernel units and associated model weights learned from the healthy source population were transferred to initialize Q-KTD training for each stroke participant, following the transfer strategy used in previous Q-KTD work (34). The resulting increase in first-epoch performance suggests that healthy-source information can place the decoder in a more informative initial region of the Q-function for some target participants.

The current results also differ meaningfully from the previous Q-KTD transfer-learning study. McDorman et al. reported improved first-epoch performance in all evaluated transfer conditions and statistically significant improvement in 23 of 24 cases when transferring between sessions from a small freewill EEG dataset (34). The larger and more heterogeneous effect observed here is expected because the current transfer problem crosses both population and dataset boundaries. The healthy and stroke datasets differed in neurological status, acquisition system, reference configuration, sampling rate, task timing, imagery content, and visual presentation. In addition, stroke-related changes may alter the neural representation of motor imagery in ways that are not present in cross-session transfer among healthy recordings. The lower consistency of benefit in the present study therefore suggests that transfer becomes less reliable as source–target domain mismatch increases.

The participant-level results illustrate this issue directly. Several participants showed gains greater than 14%, indicating that the healthy-source model provided a substantially better initialization for those cases. Conversely, other participants showed marked negative transfer, with reductions as large as 14%. The paired t-tests across the 10 Monte Carlo repetitions identified significant transfer-related differences in selected participants at p < 0.01. These tests indicate that the observed effects were consistent across repeated randomized algorithmic runs for those participants. However, the Monte Carlo repetitions represent repeated computational realizations rather than independent biological samples; therefore, the participant-level significance tests should not be interpreted as population-level clinical inference.

Although transfer learning improved mean first-epoch performance, the absolute transferred success rate of 51.82% remained only slightly above the 50% chance level for the binary decoding task. The present effect should therefore be interpreted as algorithmically meaningful but practically preliminary. The result demonstrates that source-population initialization can influence early Q-KTD learning in stroke participants, but it does not establish immediately usable BMI control. Further improvements may be obtained through optimization of kernel selection, quantization, feature normalization, exploration policy, reward design, and source-model construction. Evaluating later-epoch learning curves will also be important for determining whether transfer changes only initial performance or additionally affects convergence rate and final decoder performance.

From an assistive-technology perspective, the main value of healthy-source transfer is its potential to reduce dependence on large amounts of participant-specific stroke EEG. Stroke datasets are generally more difficult to acquire than healthy datasets because of recruitment constraints, clinical heterogeneity, fatigue, impaired motor imagery ability, and limited tolerance for lengthy calibration sessions. Reusing shared information from a comparatively large healthy source population may therefore provide a more informed starting point for individual stroke decoders. The current study did not directly measure calibration duration or the number of target-participant trials required to reach a usable threshold. Accordingly, the findings support the hypothesis that early performance gains may reduce calibration burden, but they do not demonstrate an actual reduction in calibration time.

The physiological and decoder findings should also be connected cautiously. Greater stroke-group variability, diffuse mu modulation, and altered hemispheric asymmetry provide plausible sources of mismatch between the healthy source model and individual stroke targets. However, the present analysis did not directly test whether ERD/ERS, asymmetry, band power, or time–frequency characteristics predicted transfer gain. It therefore cannot be concluded that a specific physiological feature caused positive or negative transfer. A direct feature-to-performance analysis will be needed to determine whether physiological measures can be used as transfer-selection biomarkers. Such an analysis could evaluate whether similarity to the healthy source distribution predicts Q-KTD gain or whether lesion characteristics and clinical motor impairment provide additional explanatory value.

Several methodological limitations should be considered. First, the healthy and stroke datasets were collected independently using different acquisition systems, reference configurations, task structures, imagined movements, and visual stimuli. The healthy baseline was defined immediately before motor imagery onset, whereas the stroke baseline was taken from the late-trial rest period according to the original task design. These differences may influence band-power, ERD/ERS, and baseline-normalized time–frequency comparisons independently of neurological status. Second, preprocessing did not include independent component analysis, EOG regression, systematic bad-channel rejection, or trial-level artifact rejection beyond frequency filtering. High-frequency activity, frontal channels, and unusually large participant-level values should therefore be interpreted cautiously.

Additional limitations concern temporal resolution and clinical heterogeneity. The 0.1–4 Hz band was retained to capture slow cortical and low-frequency activity, but the 0.5 s and 1.0 s analysis intervals are too short to characterize the lowest frequencies as conventional resting-state spectral slowing. These features should instead be interpreted as short-window low-frequency amplitudes. Lesion side, lesion location, impairment severity, and motor imagery ability were not incorporated into the current statistical analysis, although these factors may contribute substantially to the observed variability. The representative ERP plots were also included for descriptive illustration and were not selected using a formal group-level criterion.

Finally, the study used prerecorded EEG and a simulated center-out task rather than a closed-loop human-in-the-loop BMI. First-epoch performance provides an approximation of initial decoder behavior because each trial has not yet been repeatedly presented to the learning algorithm, but it does not reproduce user adaptation, feedback processing, fatigue, or neural nonstationarity during real-time control. Consequently, the present results demonstrate offline feasibility rather than clinical effectiveness or restored motor function.

Future work should evaluate healthy-to-stroke Q-KTD transfer in real-time closed-loop experiments and determine whether source initialization reduces the number of calibration trials or time required to reach a predefined control threshold. Additional source-domain strategies should include stroke-to-stroke transfer, in which a model trained on other stroke participants is transferred to a held-out target, and combined healthy-plus-stroke transfer, which may integrate broad healthy motor imagery structure with stroke-specific variability. Subject-specific source selection, partial transfer of selected kernel units, and early detection of negative transfer may further improve reliability. Future studies should also examine whether lesion characteristics, clinical impairment, and physiological EEG similarity can predict which source model is most appropriate for each participant.

Overall, the findings support healthy-source Q-KTD initialization as a feasible strategy for improving early offline motor imagery decoding in some individuals with acute stroke. At the same time, the near-chance average first-epoch performance and frequent negative transfer demonstrate that population-based initialization should not be applied uniformly. Reliable stroke BMI implementation will require improved Q-KTD optimization, individualized transfer selection, and prospective validation in closed-loop assistive-control settings.

## Conclusion

This study evaluated healthy-to-stroke translation for EEG-based motor imagery brain–machine interfaces by combining physiological EEG characterization with Q-KTD decoder evaluation. Healthy and acute stroke participants demonstrated shared motor imagery–related EEG activity, including post-onset mu-band modulation, while the stroke cohort showed greater interparticipant variability, more diffuse temporal patterns, and altered hemispheric asymmetry. These findings indicate that motor imagery–related neural structure is partially preserved after stroke but expressed with greater subject-specific variation.

Healthy-source transfer learning increased mean first-epoch Q-KTD success rate from 49.46% to 51.82% and improved performance in 29 of 50 stroke participants. The larger gains observed in several individuals demonstrate that information learned from a healthy EEG population can provide a beneficial initialization for Q-KTD decoding in stroke. At the same time, variability in transfer benefit indicates that the usefulness of healthy-source information depends on the correspondence between the source representation and the EEG characteristics of the target participant.

Overall, the findings support the offline feasibility of population-informed Q-KTD decoding for motor imagery BMIs after stroke. By improving early decoder performance, healthy-source transfer may reduce reliance on extensive participant-specific stroke data and potentially decrease calibration burden. Future development should focus on improving Q-KTD learning efficiency, selecting transfer sources according to individual EEG characteristics, and evaluating healthy-to-stroke, stroke-to-stroke, and combined-source transfer strategies in real-time closed-loop BMI systems.

## Acknowledgments

The authors thank the University of Kentucky Department of Electrical and Computer Engineering and Department of Biomedical Engineering for academic support during this work.

## Declaration of Interest Statement

The authors report there are no competing interests to declare.

## Funding

The authors received no financial support for the research, authorship, or publication of this article.

## Data Availability Statement

The healthy EEG data analyzed in this study are publicly available through the PhysioNet EEG Motor Movement/Imagery Dataset, version 1.0.0 (doi:10.13026/C28G6P).

The acute stroke EEG data analyzed in this study are publicly available through the Figshare repository “EEG datasets of stroke patients,” version 5, associated with Liu et al. The stroke dataset is also described in the accompanying Scientific Data article (doi:10.6084/m9.figshare.21679035.v5).

## Code Availability Statement

Custom MATLAB code used for EEG preprocessing, feature extraction, statistical analysis, and Q-KTD transfer-learning analysis will be available upon publication at the authors’ GitHub pages.

## Notes

### Competing Interest Statement

The authors have declared no competing interest.

## References

1. Wolpaw JR, Birbaumer N, McFarland DJ, Pfurtscheller G, Vaughan TM. Brain-computer interfaces for communication and control. Clin Neurophysiol. 2002;113(6):767–791. doi:10.1016/S1388-2457(02)00057-3.

2. Abiri R, Borhani S, Sellers EW, Jiang Y, Zhao X. A comprehensive review of EEG-based brain-computer interface paradigms. J Neural Eng. 2019;16(1):011001. doi:10.1088/1741-2552/aaf12e.

3. Yuan H, He B. Brain-computer interfaces using sensorimotor rhythms: current state and future perspectives. IEEE Trans Biomed Eng. 2014;61(5):1425–1435. doi:10.1109/TBME.2014.2312397.

4. Padfield N, Zabalza J, Zhao H, Masero V, Ren J. EEG-based brain-computer interfaces using motor-imagery: techniques and challenges. Sensors (Basel). 2019;19(6):1423. doi:10.3390/s19061423.

5. Vavoulis A, Figueiredo P, Vourvopoulos A. A review of online classification performance in motor imagery-based brain-computer interfaces for stroke neurorehabilitation. Signals. 2023;4(1):73–86. doi:10.3390/signals4010004.

6. Lotte F, Bougrain L, Cichocki A, Clerc M, Congedo M, Rakotomamonjy A, et al. A review of classification algorithms for EEG-based brain-computer interfaces: a 10 year update. J Neural Eng. 2018;15(3):031005. doi:10.1088/1741-2552/aab2f2.

7. Wu D, Xu Y, Lu BL. Transfer learning for EEG-based brain-computer interfaces: a review of progress made since 2016. IEEE Trans Cogn Dev Syst. 2022;14(1):4–19. doi:10.1109/TCDS.2020.3007453.

8. Grefkes C, Ward NS. Cortical reorganization after stroke: how much and how functional? Neuroscientist. 2014;20(1):56–70. doi:10.1177/1073858413491147.

9. Dodd KC, Nair VA, Prabhakaran V. Role of the contralesional vs. ipsilesional hemisphere in stroke recovery. Front Hum Neurosci. 2017;11:469. doi:10.3389/fnhum.2017.00469.

10. Kancheva I, van der Salm SMA, Ramsey NF, Vansteensel MJ. Association between lesion location and sensorimotor rhythms in stroke: a systematic review with narrative synthesis. Neurol Sci. 2023;44(12):4263–4289. doi:10.1007/s10072-023-06982-8.

11. De Vico Fallani F, Pichiorri F, Morone G, Molinari M, Babiloni F, Cincotti F, et al. Multiscale topological properties of functional brain networks during motor imagery after stroke. Neuroimage. 2013;83:438–449. doi:10.1016/j.neuroimage.2013.06.039.

12. Kaiser V, Daly I, Pichiorri F, Mattia D, Müller-Putz GR, Neuper C. Relationship between electrical brain responses to motor imagery and motor impairment in stroke. Stroke. 2012;43(10):2735–2740. doi:10.1161/STROKEAHA.112.665489.

13. Tangwiriyasakul C, Mocioiu V, van Putten MJAM, Rutten WLC. Classification of motor imagery performance in acute stroke. J Neural Eng. 2014;11(3):036001. doi:10.1088/1741-2560/11/3/036001.

14. Finnigan S, van Putten MJAM. EEG in ischaemic stroke: quantitative EEG can uniquely inform sub-acute prognoses and clinical management. Clin Neurophysiol. 2013;124(1):10–19. doi:10.1016/j.clinph.2012.07.003.

15. Bentes C, Peralta AR, Viana P, Martins H, Morgado C, Casimiro C, et al. Quantitative EEG and functional outcome following acute ischemic stroke. Clin Neurophysiol. 2018;129(8):1680–1687. doi:10.1016/j.clinph.2018.05.021.

16. Foreman B, Claassen J. Quantitative EEG for the detection of brain ischemia. Crit Care. 2012;16(2):216. doi:10.1186/cc11230.

17. van Putten MJAM, Tavy DLJ. Continuous quantitative EEG monitoring in hemispheric stroke patients using the brain symmetry index. Stroke. 2004;35(11):2489–2492. doi:10.1161/01.STR.0000144649.49861.1d.

18. Jeannerod M. Neural simulation of action: a unifying mechanism for motor cognition. Neuroimage. 2001;14(1 Pt 2):S103–S109. doi:10.1006/nimg.2001.0832.

19. Pfurtscheller G, Lopes da Silva FH. Event-related EEG/MEG synchronization and desynchronization: basic principles. Clin Neurophysiol. 1999;110(11):1842–1857. doi:10.1016/S1388-2457(99)00141-8.

20. McFarland DJ, Miner LA, Vaughan TM, Wolpaw JR. Mu and beta rhythm topographies during motor imagery and actual movements. Brain Topogr. 2000;12(3):177–186. doi:10.1023/A:1023437823106.

21. Pfurtscheller G, Neuper C, Flotzinger D, Pregenzer M. EEG-based discrimination between imagination of right and left hand movement. Electroencephalogr Clin Neurophysiol. 1997;103(6):642–651. doi:10.1016/S0013-4694(97)00080-1.

22. Graimann B, Huggins JE, Levine SP, Pfurtscheller G. Visualization of significant ERD/ERS patterns in multichannel EEG and ECoG data. Clin Neurophysiol. 2002;113(1):43–47. doi:10.1016/S1388-2457(01)00697-6.

23. Cho H, Ahn M, Ahn S, Kwon M, Jun SC. EEG datasets for motor imagery brain computer interface. Gigascience. 2017;6(7):gix034. doi:10.1093/gigascience/gix034.

24. Wilkinson CM, Burrell JI, Kuziek JWP, Thirunavukkarasu S, Buck BH, Mathewson KE. Predicting stroke severity with a 3-min recording from the Muse portable EEG system for rapid diagnosis of stroke. Sci Rep. 2020;10(1):18465. doi:10.1038/s41598-020-75379-w.

25. Peterson W, Ramakrishnan N, Browder K, Sanossian N, Nguyen P, Fink E. Differentiating ischemic stroke patients from healthy subjects using a large-scale, retrospective EEG database and machine learning methods. J Stroke Cerebrovasc Dis. 2024;33:107714. doi:10.1016/j.jstrokecerebrovasdis.2024.107714.

26. Schmidt S, Jo HG, Wittmann M, Hinterberger T. Catching the waves: slow cortical potentials as moderator of voluntary action. Neurosci Biobehav Rev. 2016;68:639–650. doi:10.1016/j.neubiorev.2016.06.023.

27. Monto S, Palva S, Voipio J, Palva JM. Very slow EEG fluctuations predict the dynamics of stimulus detection and oscillation amplitudes in humans. J Neurosci. 2008;28(33):8268–8272. doi:10.1523/JNEUROSCI.1910-08.2008.

28. Benzy VK, Vinod AP, Subasree R, Alladi S, Raghavendra K. Motor imagery hand movement direction decoding using brain computer interface to aid stroke recovery and rehabilitation. IEEE Trans Neural Syst Rehabil Eng. 2020;28(12):3051–3062. doi:10.1109/TNSRE.2020.3039331.

29. Amo C, Boquete L, de Santiago L, Barea R, Cavaliere C. Induced gamma-band activity during actual and imaginary movements: EEG analysis. Sensors (Basel). 2020;20(6):1545. doi:10.3390/s20061545.

30. Smith MM, Weaver KE, Grabowski TJ, Rao RPN, Darvas F. Non-invasive detection of high gamma band activity during motor imagery. Front Hum Neurosci. 2014;8:817. doi:10.3389/fnhum.2014.00817.

31. Gwon D, Ahn M. Alpha and high gamma phase-amplitude coupling during motor imagery and weighted cross-frequency coupling to extract discriminative cross-frequency patterns. Neuroimage. 2021;243:118403. doi:10.1016/j.neuroimage.2021.118403.

32. Ahn M, Ahn S, Hong JH, Cho H, Kim K, Kim BS, et al. Gamma band activity associated with BCI performance: simultaneous MEG/EEG study. Front Hum Neurosci. 2013;7:848. doi:10.3389/fnhum.2013.00848.

33. Jafari M, Tao X, Barua P, Tan RS, Acharya UR. Application of transfer learning for biomedical signals: a comprehensive review of the last decade. Inf Fusion. 2025;118:102982. doi:10.1016/j.inffus.2025.102982.

34. McDorman RA, Thapa BR, Kim J, Bae J. Transfer learning in EEG-based reinforcement learning brain machine interfaces via Q-learning Kernel Temporal Differences. Paper presented at: 47th Annual International Conference of the IEEE Engineering in Medicine and Biology Society; 2025 Jul 14–18; Copenhagen, Denmark.

35. DiGiovanna J, Mahmoudi B, Fortes J, Principe JC, Sanchez JC. Coadaptive brain-machine interface via reinforcement learning. IEEE Trans Biomed Eng. 2009;56(1):54–64. doi:10.1109/TBME.2008.926699.

36. Sutton RS, Barto AG. Reinforcement learning: an introduction. Cambridge (MA): MIT Press; 1998. 37.

37. Watkins CJCH, Dayan P. Q-learning. Mach Learn. 1992;8(3-4):279–292. doi:10.1007/BF00992698.

38. Bae J, Sanchez Giraldo LG, Pohlmeyer EA, Francis JT, Sanchez JC, Principe JC. Kernel temporal differences for neural decoding. Comput Intell Neurosci. 2015;2015:481375. doi:10.1155/2015/481375.

39. Thapa BR, Tangarife DR, Bae J. Kernel temporal differences for EEG-based reinforcement learning brain machine interfaces. Paper presented at: 44th Annual International Conference of the IEEE Engineering in Medicine and Biology Society; 2022 Jul 11–15; Glasgow, UK.

40. Pareigis S. Algorithmic foundations of reinforcement learning. In: Unger H, Schaible M, editors. Advances in real-time and autonomous systems. Cham (Switzerland): Springer Nature; 2024. p. 1–27.

41. Liu W, Principe JC, Haykin S. Kernel adaptive filtering: a comprehensive introduction. Hoboken (NJ): John Wiley & Sons; 2010.

42. Chen BD, Zhao S, Zhu P, Principe J. Quantized kernel least mean square algorithm. IEEE Trans Neural Netw Learn Syst. 2012;23(1):22–32. doi:10.1109/TNNLS.2011.2178446.

43. Schalk G. EEG Motor Movement/Imagery Dataset (version 1.0.0). PhysioNet. 2009. RRID:SCR_007345. doi:10.13026/C28G6P.

44. Schalk G, McFarland DJ, Hinterberger T, Birbaumer N, Wolpaw JR. BCI2000: a general-purpose brain-computer interface (BCI) system. IEEE Trans Biomed Eng. 2004;51(6):1034–1043. doi:10.1109/TBME.2004.827072.

45. Baravalle R, Rosso O, Montani F. Causal Shannon–Fisher Characterization of Motor/Imagery Movements in EEG. Entropy. 2018;20:660. doi: 10.3390/e20090660.

46. Liu H, Wang Y, Zhang X, Li Y, Chen X, Wang C, et al. An EEG motor imagery dataset for brain computer interface in acute stroke patients. Sci Data. 2024;11(1):69. doi:10.1038/s41597-023-02787-8.

47. Liu H, Lv X. EEG datasets of stroke patients [dataset]. Figshare. 2023. doi:10.6084/m9.figshare.21679035.v5.

48. Welch BL. The generalization of “Student’s” problem when several different population variances are involved. Biometrika. 1947;34(1-2):28–35. doi:10.1093/biomet/34.1-2.28.

49. Benjamini Y, Hochberg Y. Controlling the false discovery rate: a practical and powerful approach to multiple testing. J R Stat Soc Series B Stat Methodol. 1995;57(1):289–300.

50. Cohen J. Statistical Power Analysis for the Behavioral Sciences. 2nd ed. Hillsdale (NJ): Lawrence Erlbaum Associates; 1988.

